# RAS isoform specific activities are disrupted by disease associated mutations during cell differentiation

**DOI:** 10.1101/2023.10.05.561021

**Authors:** Rohan Chippalkatti, Bianca Parisi, Farah Kouzi, Christina Laurini, Nesrine Ben Fredj, Daniel Kwaku Abankwa

## Abstract

The Ras-MAPK pathway is aberrantly regulated in cancer and developmental diseases called RASopathies. While typically the impact of Ras on the proliferation of various cancer cell lines is assessed, it is poorly established how Ras affects cellular differentiation.

Here we implement the C2C12 myoblast cell line to systematically study the effect of Ras mutants and Ras-pathway drugs on differentiation. We first provide evidence that a minor pool of Pax7+ progenitors replenishes a major pool of transit amplifying cells that are ready to differentiate. Our data indicate that Ras isoforms have distinct roles in the differentiating culture, where K-Ras is more important than N-Ras to maintain the progenitor pool and H-Ras is significant for terminal differentiation. This assay could therefore provide significant new insights into Ras biology and Ras-driven diseases.

In line with this, we found that all oncogenic Ras mutants block terminal differentiation of transit amplifying cells. Notably, while RASopathy K-Ras variants that are also NF1-GAP resistant also block differentiation, albeit less than their oncogenic counterparts. Profiling of targeted Ras-pathway drugs on oncogenic Ras mutants revealed their distinct abilities to restore normal differentiation as compared to triggering cell death. In particular, the MEK-inhibitor trametinib could broadly restore differentiation, while the mTOR-inhibitor rapamycin broadly suppressed differentiation.

We expect that this quantitative assessment of the impact of Ras-pathway mutants and drugs on cellular differentiation has great potential to complement cancer cell proliferation data.

## Introduction

Malignant tumors are characterized by abnormal proliferation and invasive growth of dedifferentiated tissue. The Ras-pathway is central to control cellular proliferation, differentiation and survival, and is dysregulated in virtually every cancer (Crespo & Leon, 2000; Hanahan, 2022). Three *RAS* genes, *KRAS*, *NRAS* and *HRAS,* are mutated in 19 % of human cancers making *RAS* the most frequently mutated oncogene (Prior *et al*, 2020). Out of the two *KRAS* splice isoforms, K-Ras4A and K-Ras4B, the latter is the highest expressed isoform and the major focus of current drug development (Hood *et al*, 2023; Moore *et al*, 2020; Tsai *et al*, 2015).

Ras membrane association is required for its activity, and membrane affinity is mediated by C-terminal lipid modifications of Ras by farnesyltransferase and palmitoyltransferases (Pavic *et al*, 2022). Farnesylation also mediates binding of Ras to trafficking chaperones, such as PDE6D and calmodulin, which facilitate its diffusion, followed by trapping on secretory organelles, and subsequent vesicular transport to the plasma membrane (Schmick *et al*, 2015).

Canonical Ras signaling emerges at the plasma membrane, where extracellular mitogens activate receptor tyrosine kinases, such as epidermal growth factor receptor (EGFR), which indirectly relays its activation to guanine nucleotide exchange factors (GEFs), such as SOS. GEFs facilitate exchange of GDP for GTP, thus activating Ras. The active GTP-bound Ras then recruits effector proteins, such as Raf, PI3K and RalGDS from the cytosol to the membrane, leading to their activation (Simanshu *et al*, 2017). Raf-kinases trigger the MAPK-pathway, which includes activation of downstream kinases MEK and ERK, the latter of which leads to well characterized changes that drive the cell cycle and thus proliferation (Crespo & Leon, 2000). The effector PI3K activates the kinase Akt, which further downstream engages the mTORC1-pathway and thus cell growth and many other crucial cellular processes (Laplante & Sabatini, 2013). Ras can furthermore activate mTORC2, which phosphorylates Akt on Ser473 (Kovalski *et al*, 2019).

The active state of Ras is tightly regulated, with GTP-Ras becoming inactivated by GTPase-activating proteins (GAPs) (Simanshu *et al*., 2017). The most prominently studied GAP is neurofibromin 1 (NF1), which is recruited aided by B-Raf and one of three SPRED proteins to K-Ras nanodomains of the plasma membrane (Siljamaki & Abankwa, 2016; Stowe *et al*, 2012; Yan *et al*, 2020). Landmark structural data from the mid 1990s already explained how hotspot oncogenic mutations in codons 12 and 61 of Ras disable the GTP-hydrolysis of Ras by NF1 and other arginine-finger GAPs (Ahmadian *et al*, 1997; Scheffzek *et al*, 1997). However, the heterotrimeric G protein GAP RGS3 with a catalytic asparagine, was recently shown to facilitate GTP-hydrolysis of all major oncogenic K-Ras mutants (G12D/V, G13C/D) (Li *et al*, 2021), suggesting that a distinct function of NF1 is disabled by oncogenic Ras.

In line with the mitogen-independent mutational activation of Ras, cancer cell assays typically assess the uncontrolled cell growth. Proliferation assays provided a wealth of data in cancer research, such as from large scale genetic and chemical screens (Barretina *et al*, 2012; McDonald *et al*, 2017; Tsherniak *et al*, 2017). However, among pathologists it is well established that dedifferentiation is the most unsettling hallmark of cancer (Chaffer & Weinberg, 2015; Hanahan, 2022). Unfortunately, the functions of Ras during cellular differentiation are only poorly understood and typically not assayed.

This lack of understanding also impacts on the treatment development for another type of Ras-driven diseases. Germline mutations in the RAS-MAPK pathway lead to individually rare but collectively common developmental syndromes called RASopathies, which are characterized by facial malformations, short stature, cutaneous defects, cardiac hypertrophy and a pre-disposition to cancer (Castel *et al*, 2020; Rauen, 2013). They illustrate how even a mild overactivation of the MAPK-pathway during embryonal development perturbs proper differentiation in multiple organ systems.

For instance, the RASopathy Noonan syndrome can be caused by the *KRAS-D153V* mutation, which in contrast to cancer-associated hotspot mutations, is still sensitive to the GAP NF1, but shows mildly increased effector binding (Gremer *et al*, 2011). Loss-of-function mutations in *NF1* itself lead to neurofibromatosis type I, one of the more common RASopathies (Rauen, 2013). This disorder shares some phenotypic similarities with the very rare RASopathy Legius syndrome, which is caused by heterozygous loss-of-function mutations in the *SPRED1* gene (Brems *et al*, 2012). To analyze RASopathy mutants, dedicated low-throughput assays have been developed, which characterize early developmental defects during gastrulation or later in the whole organism in zebrafish and mouse animal models (Jindal *et al*, 2015). Yet, insufficient developmental or cell differentiation assay capacities may underlie the lack of efficacious therapies for most RASopathies (Gross *et al*, 2020).

These observed developmental defects in RASopathies are consistent with the deep integration of MAPK-signaling already at the level of stem cell maintenance. During organismal development, pluripotent stem cells give rise to a vast variety of differentiated tissues (Morrison & Kimble, 2006). Both during priming of naïve mouse embryonic stem cells and maintenance of pluripotency in human induced pluripotent stem cells is the MAPK-pathway involved (Altshuler *et al*, 2018; Haghighi *et al*, 2018). In the fully developed organism, adult stem and progenitor cells are important for tissue homeostasis. They typically divide asymmetrically, giving rise to one stem cell (referred to as self-renewal) and one committed or differentiated cell (Morrison & Kimble, 2006).

The C2C12 cell model is one of the best characterized *in vitro* differentiation systems, which recapitulates essential *in vivo* processes (Yin *et al*, 2013). This cell line was derived from a skeletal muscle of a 2-month old mouse, and is typically considered a heterogenous population of myoblasts (myogenic progenitor cells), which proliferate and remain undifferentiated under high serum conditions (Bennett & Tonks, 1997). Mitogen withdrawal in low serum culture conditions, rapidly triggers terminal differentiation of the majority of C2C12 cells into multinucleated myotubes within five days (Bennett & Tonks, 1997).

However, it is known that a small fraction of proliferating C2C12 cells expresses the muscle progenitor marker Pax7, a paired box transcription factor, as well as the basic helix-loop-helix transcription factor Myf5 (Yoshida *et al*, 1998). Differentiating cells downregulate Pax7 and upregulate myogenic factors, such as MyoD, myogenin and subsequently late differentiation markers, such as the motor protein myosin II heavy chain (MyHC) (Brown *et al*, 2012; Olguin & Olwin, 2004). Interestingly, upon serum withdrawal a minor fraction remains undifferentiated and continues to express the progenitor marker Pax7, but no MyoD, Myf-5 or myogenin (Olguin & Olwin, 2004; Yoshida *et al*., 1998). These features strikingly resemble quiescent satellite cells, the Pax7 positive adult stem cell population in muscles (Yablonka-Reuveni & Rivera, 1994). Currently, it is not fully resolved, how the minor fraction of myoblast progenitors is connected to a major fraction of differentiated cells.

Myogenic differentiation is initiated by a rapid upregulation of SPRED1 upon serum switching and a subsequent decrease in MAPK-signaling (Bennett & Tonks, 1997; Wakioka *et al*, 2001). In line with mitogens maintaining proliferation of myoblasts, oncogenic Ras prevents myogenic differentiation by downregulating the myogenic transcription factor MyoD and myogenin (Lassar *et al*, 1989). Conversely, overexpression of tumor suppressors, such as Sprouty2 and SPRED1 stimulate myogenesis even under high serum conditions (de Alvaro *et al*, 2005; Wakioka *et al*., 2001). Terminal differentiation is then promoted by mTORC2-Akt activity (Shu & Houghton, 2009).

The proper differentiation trajectories of tissues is perturbed in cancer and may lead to the emergence of rare cancer stem cells, which alone have the potential to seed new tumors, for example during metastization and relapse after therapy (Morrison & Kimble, 2006). Current cancer stem cell models suggest either reprogramming of differentiated cells or an evolution directly from transformed stem/ progenitor cells (Ansieau, 2013; Batlle & Clevers, 2017). Cancer stem cells are best characterized in functional and lineage tracing assays *in vivo* (Nassar & Blanpain, 2016). Yet, *in vitro* surrogate assays persist, such as flow cytometry based detection of cancer stem cell markers (e.g., CD44+/CD24-), or of cancer stem cells in the side population, which is characterized by their increased drug efflux properties (Golebiewska *et al*, 2011; Li *et al*, 2017). In addition, low serum, non-adherent 3D spheroid cultures of human mammary stem/ progenitors cells, called mammospheres, were originally employed to maintain and study such cells in culture (Dontu *et al*, 2003). Subsequently these culture conditions were widely adopted to monitor cancer cell stemness from tumorospheres (Weiswald *et al*, 2015). The simplicity of this assay enabled screening for compounds that may have a potential to target specifically cancer stem cells (2022; Mathews *et al*, 2012; She *et al*, 2021).

Current data suggest that *KRAS* is the strongest driver of stemness features, followed by *NRAS* and *HRAS* (Najumudeen *et al*, 2016; Quinlan *et al*, 2008; Wang *et al*, 2015). This potency order that was obtained across multiple model systems strikingly correlates with the *RAS* mutation frequency in cancer (Chippalkatti & Abankwa, 2021). Salinomycin was one of the first cancer stem cell selective inhibitors that was described by the Weinberg group (Gupta *et al*, 2009). This and related compounds showed selective activity against K-Ras, but not H-Ras, suggesting that K-Ras is of high significance in cancer stem cells (Najumudeen *et al*., 2016; Okutachi *et al*, 2021; Siddiqui *et al*, 2021). While these natural products may become starting points for the development of more potent drugs, they contrast with dedicated inhibitors raised against the target K-Ras. Two such inhibitors, sotorasib (AMG 510) and adagrasib (MRTX849) have recently been approved (Canon *et al*, 2019; Fell *et al*, 2020). Essentially all of these covalent K-Ras-G12C specific inhibitors were built on the development of the compound ARS- 1620 (Janes *et al*, 2018). Moreover, MRTX849 derivatives gave rise to other non-covalent inhibitors, including MRTX1133 against K-Ras-G12D (Wang *et al*, 2022). While allele specific inhibitors promise uniquely small side-effects, their limited applicability necessitates the development of innovative K-Ras inhibitors with new modes of action (Steffen *et al*, 2023).

Here we utilized mainly flow cytometry-based differentiation marker quantification to understand the impact of the three Ras isoforms, notably K-Ras4B, on C2C12 cell differentiation. We first establish a new baseline of understanding C2C12 cell differentiation, by elaborating that Pax7+ progenitors replenish a major pool of Pax7-transit amplifying cells, which then give rise to the MyHC+ differentiated cells.

We then elaborate the distinct impact of Ras isoforms on differentiation using specific genetic perturbations. Finally, we demonstrate the applicability of our assay for medium throughput assessment of Ras-pathway drugs to restore differentiation that was perturbed by disease-associated Ras-isoforms and -alleles. Our results demonstrate how profoundly Ras-isoforms impact on cell differentiation and demonstrate how to rapidly analyze the effect of Ras-mutants and -drugs on differentiation.

## STAR Methods

### Key resources table

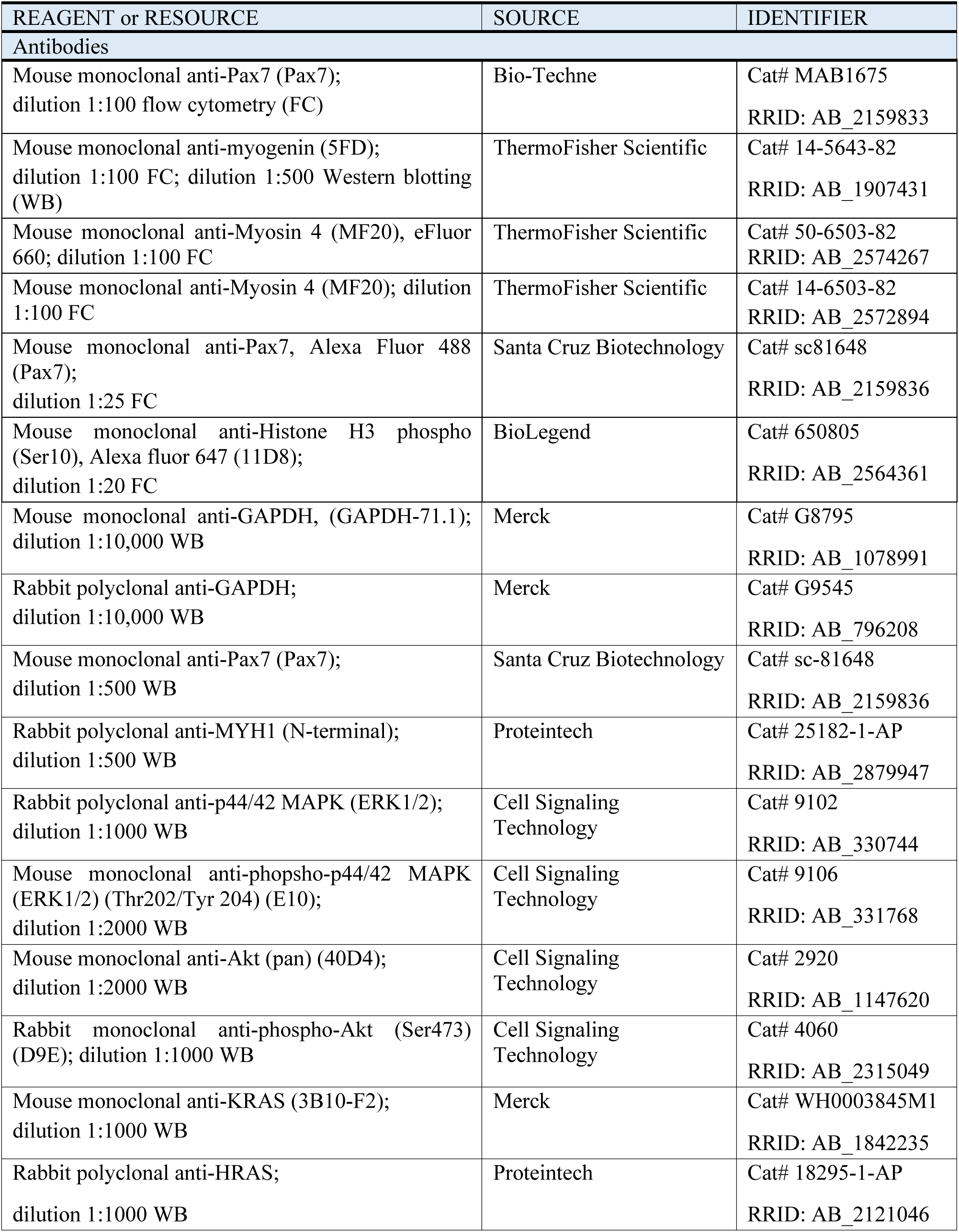
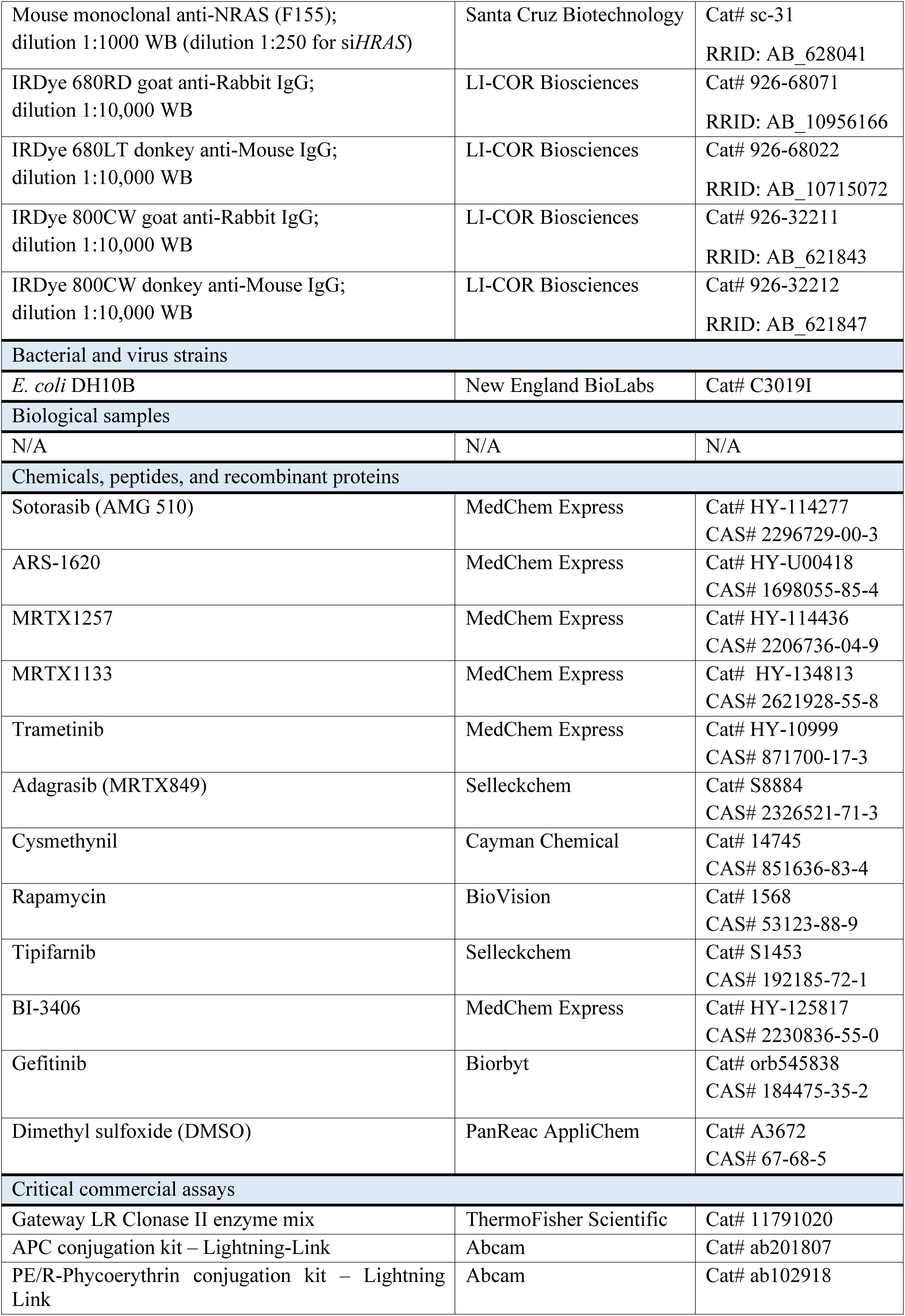
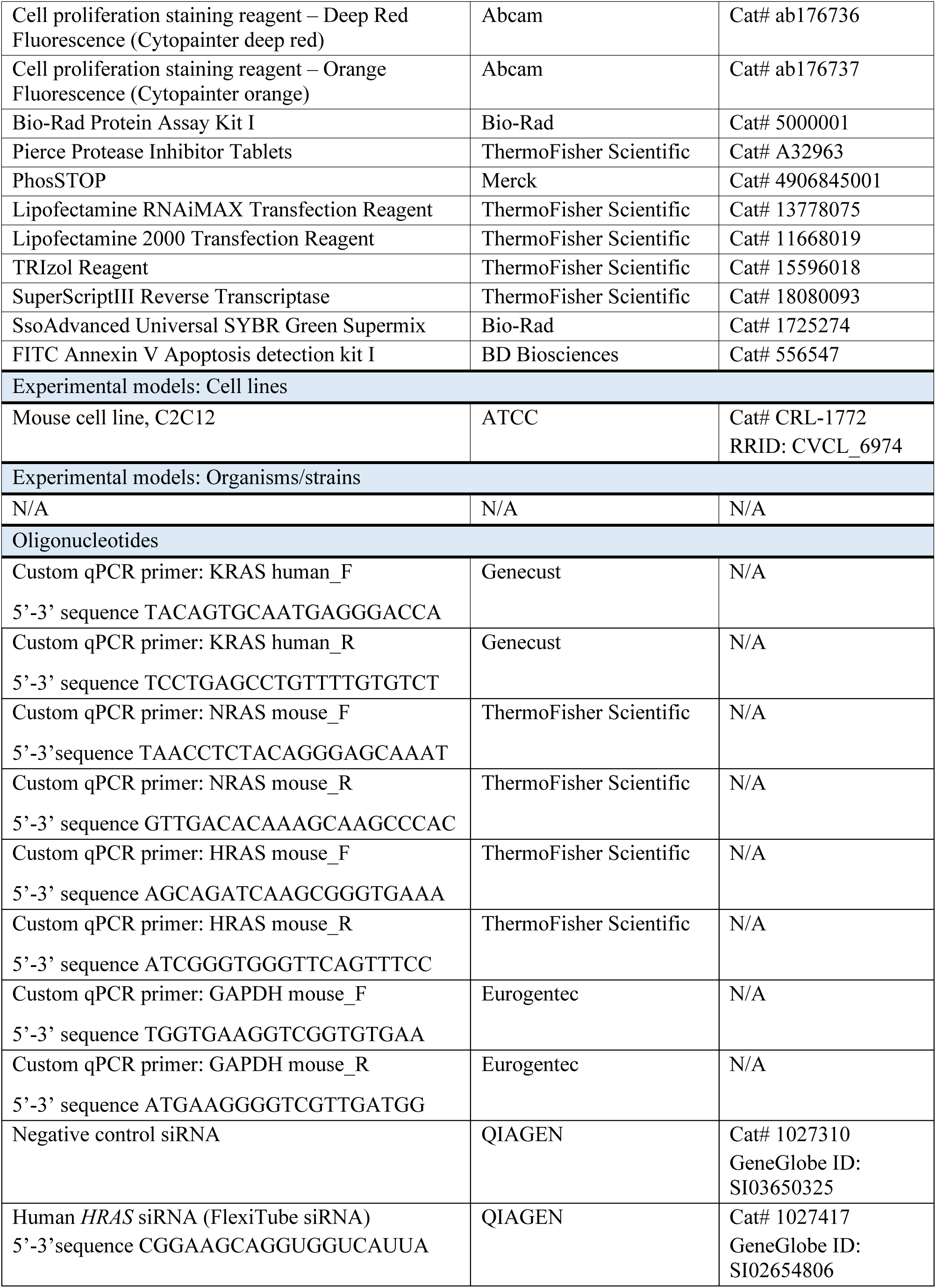
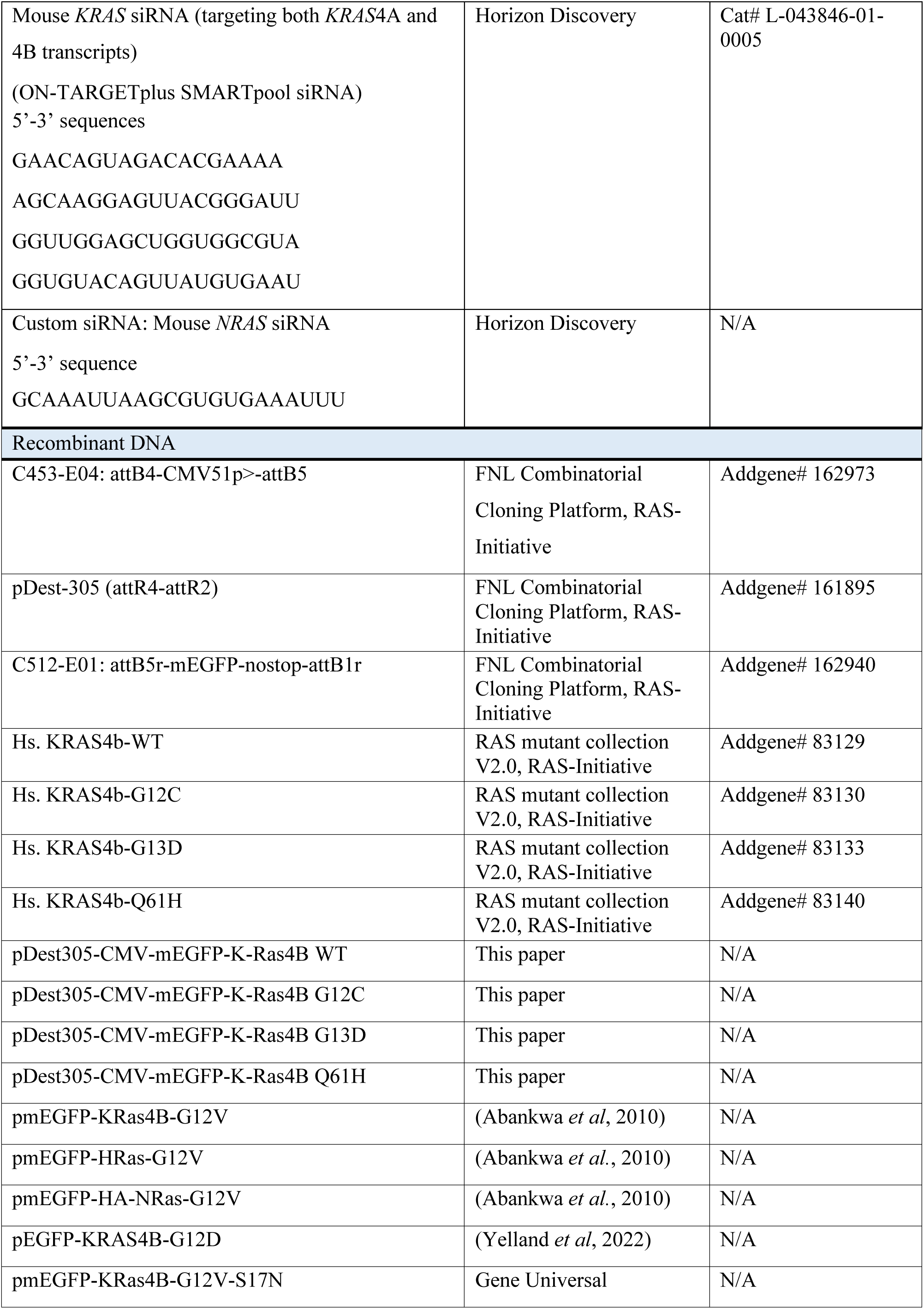
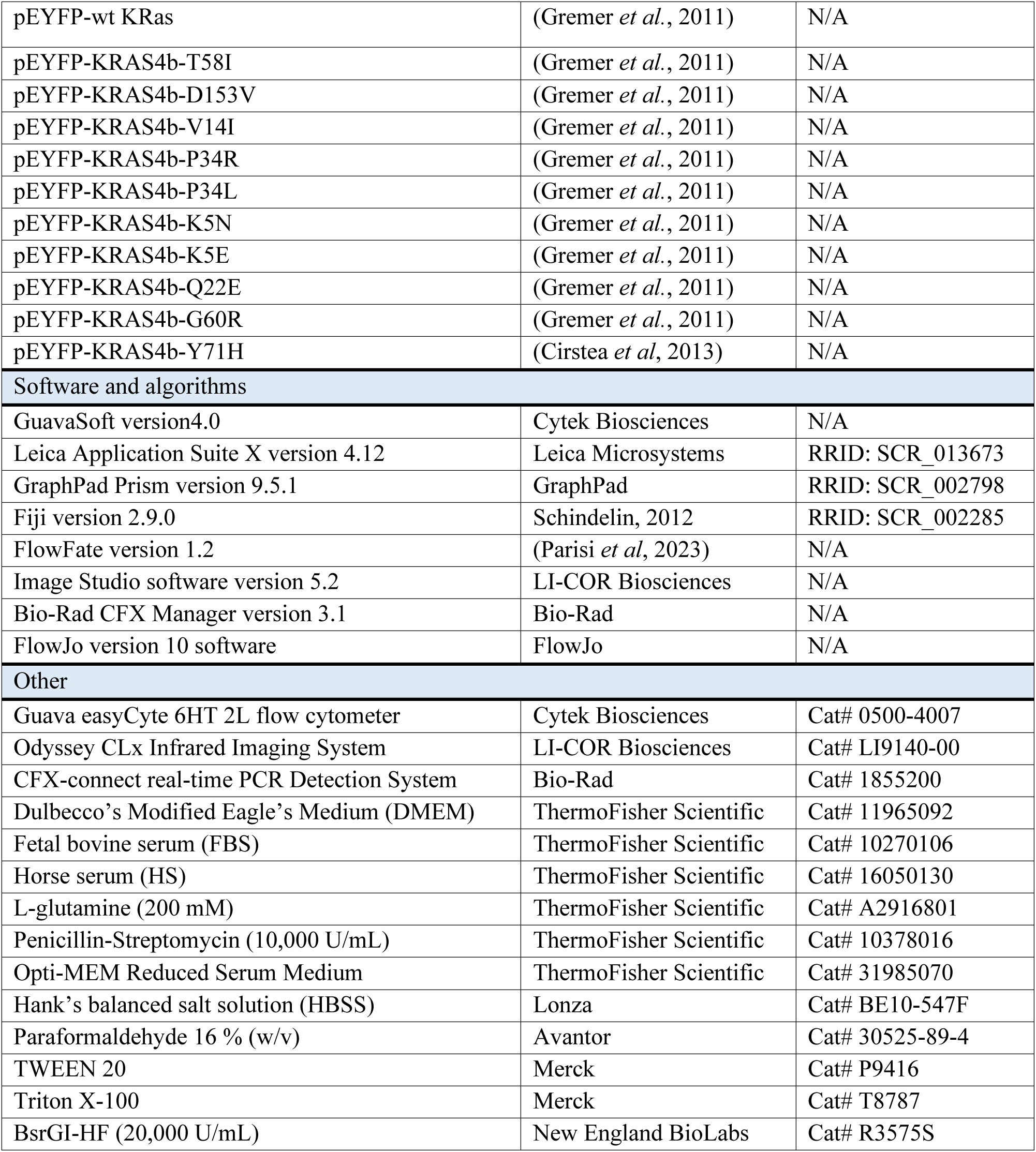

### Cell Culture and transfections

C2C12 cells were maintained in Dulbecco’s modified Eagle Medium (DMEM) supplemented with ∼9 % (v/v) fetal bovine serum (FBS), 2 mM L-glutamine and 1 % penicillin/streptomycin (high serum culture medium). Cells were incubated in a humidified incubator at 37 °C, with 5 % CO_2_ and passaged when reaching 50 – 60 % confluency. Passage numbers are indicated in the legends and passages beyond ten were not employed. Cell culture medium was exchanged with DMEM supplemented with ∼2 % horse serum (HS), 2 mM L-glutamine and 1 % penicillin/streptomycin (low serum culture medium) to induce differentiation at 90 % confluency. For experiments lasting three to five days fresh DMEM + 2% HS was added every day.

### Preparation of expression constructs

Plasmid constructs used for transfection were generated by multisite gateway cloning (Wall *et al*, 2014). To generate the final expression constructs, LR Clonase II enzyme mix was used to perform recombination of the entry clones encoding the promoter, the fluorescent tag and the gene of interest with the pDest-305 destination vector. The reaction mix was transformed into ccdB-sensitive *E. coli* strain DH10B. Colonies resistant to ampicillin were selected for screening of their plasmid DNA by restriction digestion with BsrGI-HF enzyme. Plasmids with the expected fragment sizes were then further validated by sequencing. The pmEGFP-KRas4B-G12V-S17N construct was generated by site directed mutagenesis on the pmEGFP-KRas4B-G12V background by Gene Universal (Delaware, USA).

### Plasmid DNA transfection

C2C12 cells were seeded in high serum medium at a density of 100,000 cells per mL of a 6- well plate (#10062-892, Avantor) and transfection was performed after the cells reached 50 – 60 % confluency (typically 24 h after seeding). Transfection was performed according to the protocols provided by the manufacturer. 2 µg of plasmid DNA and 7.5 µL Lipofectamine 2000 were added to 250 µL Opti-MEM medium. The mixture was vortexed, incubated at 22 - 25 °C for 10 min and subsequently added dropwise to one well of a 6-well plate. Medium was exchanged with fresh high serum medium after 4 h. After further 24 h incubation with high serum, medium was exchanged with low serum medium to induce differentiation.

### Knockdown by siRNA transfection

Any siRNAs were transfected using Lipofectamine RNAiMAX diluted in Opti-MEM medium. Typically, 100 nM of each siRNA and 7.5 µL of Lipofectamine were added to 250 µL of Opti-MEM medium, mixed and incubated at 22 – 25 °C for 10 min. Cells were transfected at 50 – 60 % confluency by adding this mixture. After 24 h of incubation with the siRNA-Lipofectamine mixture, the medium was substituted with fresh high serum medium. After 3 – 4 h, high serum medium was exchanged with low serum medium, and cells were cultured in low serum medium that was replaced every day.

### Antibody conjugation with Allophycocyanin (APC) or Phycoerythrin (PE)

For flow cytometry experiments anti-Pax7 and anti-myogenin antibodies were conjugated to APC with the reagents provided in the APC conjugation kit – Lightning link. Anti-Myosin 4 Monoclonal antibody (anti-MyHC) was conjugated with PE with the components provided in the PE/R-Phycoerythrin conjugation kit – Lightning link. The modifier reagent and quencher reagent were contained in both kits. Additionally, the APC conjugation reaction mix was supplied with the APC kit and the PE conjugation reaction mix was supplied with the PE/R-Phycoerythrin kit. Anti-Pax7 or anti-myogenin antibodies at a concentration of 1 mg/ mL and 100 μL volume were mixed with 100 μg APC Conjugation Reaction Mix, together with 10 μL of Modifier Reagent. Similarly, 100 μL of 0.5 mg/mL anti-MyHC antibody was mixed with 100 μg PE Conjugation Reaction Mix with 10 μL of Modifier Reagent. This mixture was stored in the dark at 22 – 25 °C for 3 – 4 h. Subsequently, the conjugation reaction was stopped by adding 10 μL quencher reagent to the conjugation mix. The conjugated antibody solution was used at a dilution of 1:100 and stored at 4 °C for up to 6 months.

The degree of labelling (DOL) was calculated according to the formula, DOL = (Amax × MW × dilution factor)/(ε × [conjugate]). MW is the molecular weight of IgG (∼150,000 g/mol), and ε is the molar extinction coefficient of the dye (e.g. 700,000 M^-1^ cm^-1^ for APC), [conjugate] = {[A280 – (Amax × Cf)]/1.4} × dilution factor. Here, [conjugate] is the concentration in mg/ mL of the antibody conjugate, ‘dilution factor’ is the fold of dilution used for spectral measurements. A280 is the absorbance of the conjugate at 280 nm and Amax is the absorption maximum of APC. Cf is the absorbance correction factor and the value 1.4 is the extinction coefficient of IgG in mg/mL. For APC, Cf is 0.22. We obtained 0.5 – 1 moles of APC per antibody, which falls within the optimal range for labelling. Similar results were obtained with the PE conjugation kit.

### Flow cytometry

Cells were seeded at a concentration of 100,000 cells/mL per well of a 6-well plate. This was followed by transfection with the required plasmids or siRNAs and a low serum culture period for the time indicated in the figures or figure legends. Fresh low serum, where required with drug/ compound, was added every day. Cells were then harvested by trypsinization with 0.05 % (w/v) trypsin EDTA (1×) for 5 min and pelleted by centrifugation at 500 × g for 5 min. The subsequent steps were performed at 22 - 25 °C. The cell pellet was subsequently fixed with 4 % (w/v) paraformaldehyde (PFA) in PBS for 10 min. After washing with PBS, cells were permeabilized with 0.5 % (v/v) Triton X-100 in PBS for 10 min. Subsequently cells were washed with 0.05 % (v/v) Tween 20 in PBS (PBST) and immunolabelled with fluorescent dye-conjugated primary antibodies at the indicated dilutions in PBST for 1 h at 4 °C. Subsequently cells were pelleted by centrifugation at 500 × g for 5 min and resuspended in PBST for flow cytometric analysis.

We first gated around 30,000 cells based on their forward and side scatter properties to exclude debris or dead cells. The intact cells thus obtained were further analyzed for expression markers using dot plots. In general, 1,000 to 10,000 EGFP-variant construct expressing intact cells were collected, depending on the treatment condition. Predominantly, mEGFP-tagged constructs were employed, which have the identical brightness as EGFP-tagged constructs (Norris *et al*, 2015). EGFP-variants and Alexa Fluor 488 were excited with the 50 mW photodiode 488 nm laser and the fluorescent signal was detected using the Green-B filter (band pass 525/30 nm) on a Guava Luminex easyCyte 6HT 2L flow cytometer. APC or eFluor 660 were excited using the 100 mW photodiode 642 nm laser and detected using the Red-R filter (band pass 661/15 nm). PE was excited using the 50 mW photodiode 488 nm laser and detected using the Yel-B filter (band pass 583/26 nm). An unlabeled control sample was always included as a control to correct for autofluorescence. Spectral overlap of PE and EGFP was corrected via digital compensation using single color EGFP-only and PE-only control samples.

Labelled or EGFP-variant expressing cells were quantified using the ‘Quad Stat plot’ feature on the GuavaSoft 4.0 software. The laser power and the voltage gain settings were adjusted such that the unlabeled events appeared below the level of 10^1^ relative fluorescence units (RFU). Two EGFP-expression windows were defined, ‘GFP low’ between 10^1^ and 10^2^ RFU and ‘GFP high’ between 10^2^ and 10^5^ RFU. This allowed to quantitate differentiation in dependence of the EGFP-expression level. The majority of data were analyzed in the GFP low window, which produced a phenotype comparable to non-transfected C2C12 cells. A detailed step-by-step procedure of sample preparation, data acquisition and analysis is described in our previous work (Parisi *et al*., 2023).

To visualize the changes in Pax7+, myogenin+ and MyHC+ cell fractions over time, histograms depicting the fluorescence intensities of marker-labelled populations were generated with the Overlays tool in FlowJo.

To evaluate drug-induced cell toxicity, gating was applied based on forward and side scatter characteristics to separate total events into intact cells and debris. The percentage of intact cells among the total events was then plotted for each EGFP-construct and drug treatment to estimate cell toxicity.

### Cell proliferation analysis in dye dilution experiments

A 500 × stock solution of Cytopainter deep red/Cytopainter orange was prepared as described by the manufacturer and stored at -20°C. A 1 × working solution was prepared on the day of labelling in Hank’s balanced salt solution (HBSS). Cells from a confluent flask were detached with trypsinization, pelleted by centrifugation at 200 × *g* for 3 min and 1 mL high serum medium was added to the pellet. Cells were then counted on a Z1 particle counter (Beckman Coulter) to obtain 100,000 cells per 1 mL high serum medium. Cells were again pelleted by centrifugation at 500 × *g* for 5 min, resuspended in 500 μL cytopainter working solution and incubated in the dark at 37 °C for 15 min. Cells were once more pelleted (500 × *g* for 5 min) and the pellet was washed once with 500 μL HBSS. After another round of pelleting, cells were finally resuspended in 1 mL high serum medium. 200 μL from this cell suspension was kept aside as the day 0 followed by fixation with 4 % (w/v) PFA. Out of the remaining 800 μL, 200 μL cell suspension containing 20,000 cells was dispensed per well in 3 wells of a 6-well plate containing 2 mL high serum medium. Cells were then incubated for 3 days in the cell culture incubator. Cells from one well were collected by trypsinization on day 1, day 2 and day 3 and processed for immunofluorescence with mouse monoclonal anti-Pax7, Alexa Fluor 488 antibody. Cytopainter deep red fluorescence was detected by 642 nm laser excitation and using the Red-R emission filter (band pass 661/15 nm). The half-life (t_1/2_) of the cytopainter dilution was calculated using the one-phase decay equation in Prism 9.0, Y = (Y0 – plateau) × exp(-K × X) + plateau. Here, X is time (days), Y is the cytopainter mean fluorescence intensity, Y0 is the Y value at day 0 which decays with one phase to plateau and K is the rate constant expressed as a reciprocal of the X axis unit. The t_1/2_ value was calculated as ln(2)/K.

The ‘Misc parameters’ feature on the GuavaSoft software provides the number of events per µL of each sample, which allowed us to calculate the cell concentration per mL of Pax7+ and Pax7-fractions.

To analyze cell proliferation in cells transfected with mEGFP-tagged wt K-Ras or K-RasG12V constructs, 100,000 cells/ mL were first seeded in all wells of a 6-well plate. Each construct was transfected in all wells of a 6-well plate. Transfected cells were then harvested by trypsinization, cells from all wells were pooled in a single 15 ml falcon tube and cell concentration was determined with a Z1 particle counter. Cell staining with cytopainter orange reagent was then carried out in exactly as described above for cytopainter deep red. Cells labelled with cytopainter orange were then fixed and immunolabelled with APC conjugated anti-Pax7 antibody. Cytopainter orange fluorescence was detected with 488 nm laser excitation and the Yel-B emission filter (band pass 583/26 nm). Cytopainter orange fluorescence signal for Pax7+ and Pax7-fractions was measured specifically for cells in the ‘GFP low’ window.

### Cell toxicity analysis with 7-AAD

The cell impermeable dye 7-Aminoactinomycin D (7-AAD) is excluded from intact cells but intercalates into the genomic DNA of late apoptotic and necrotic cells since the plasma membrane integrity of these cells is compromised. The toxicity of K-RasG12C inhibitors was analyzed in cell fractions transfected with pDest305-CMV-mEGFP-K-RasG12C. Cells were treated 24 h after transfection with DMSO, sotorasib (AMG 510), MRTX1257 and adagrasib (MTRX849) in low serum for three days at indicated concentrations. The medium was first aspirated from drug treated cells to collect the ‘supernatant’ fraction of dead cells. The ‘adherent’ fraction of cells was collected by trypsinization. After centrifugation at 500 × *g* for 5 min and a washing step with PBS, both fractions were labelled with the provided 7-AAD solution for 15 min on ice according to the manufacturer instructions. The 7-AAD fluorescence was detected using the 100 mW photodiode 642 nm laser and the Red-B filter (band pass 635/40 nm) by flow cytometry. Cells expressing EGFP-constructs and labelled with 7-AAD were gated and counted using the Quad Stat plot feature on GuavaSoft software. Percentages of GFP+ 7AAD+ cells were quantified from the GFP low window and plotted for each drug treatment.

### Microscopy

Brightfield images were acquired with 10 and 20 × objectives of a Leica DMI3000 B inverted microscope equipped with a Leica DFC360 FX digital camera. Samples were illuminated using pE400max LED white light source from CoolLED. Images were analyzed with Leica LAS X software.

### Immunoblotting

Cells were seeded at a density of 100,000 cells/ mL in each well of a 6-well plate, transfected and then cultured in low serum as indicated in the figures. Cells were then washed with ice-cold PBS and lysed for 30 min on ice using a total of 100 μL of lysis buffer (10 mM Tris pH 7.5, 150 mM NaCl, 0.5 mM EDTA, 0.2% NP40) supplemented with one tablet of protease inhibitor cocktail per 10 mL lysis buffer. The lysis buffer was additionally supplemented with one tablet per 10 mL of the PhosSTOP phosphatase inhibitor cocktail. Cells were collected using a scraper and incubated on ice for 30 min with intermittent vortexing. Lysates were cleared by centrifugation at 13,000 × *g* for 10 min at 4 °C. Supernatants were collected and quantified by the Bradford assay using Bio-Rad Protein Assay Kit, followed by heating at 95 °C for 5 min. Protein samples were then resolved on denaturing SDS-PAGE. A 6 % resolving gel was used for detection of MyHC, 10 % resolving gel for Pax7, ERK1/2, Akt and 15 % resolving gel for detection of myogenin, K-Ras (total K-Ras, both 4A and 4B isoforms), H-Ras, N-Ras. Subsequently, separated proteins were transferred onto a nitrocellulose blotting membrane 0.2 μm by using a TransBlot turbo Transfer System (Bio-Rad). The blots were probed as indicated in the figures with the antibodies diluted as described in the Key Resources Table. The membranes were washed 3 times with PBST (0.2 % (v/v) Tween 20 in PBS) and then incubated with anti-mouse or anti-rabbit IRDye800CW or IRDye680RD/LT conjugated secondary antibodies. Finally, protein bands were detected using a LI-COR ODYSSEY CLx system. Band intensities were quantified using Fiji and normalized to GAPDH. The relative abundances of phosphorylated ERK1/2 and phospho-Akt were quantified as the ratios of the intensities of phosphorylated proteins and the total proteins.

### Quantitative RT-PCR of gene transcripts

Cells were seeded at a density of 100,000 cells/ mL of a 6-well plate and transfected with siRNAs directed against *KRAS*, *NRAS* and *HRAS*. Note that the *HRAS* siRNA against the human mRNA was also targeting the identical sequence of the mouse mRNA.

After culturing in low serum medium, cells were collected as indicated in the figures. Total RNA was isolated using Trizol according to the manufacturer’s protocol. Reverse transcription was performed with 1 µg of total RNA using SuperScriptIII Reverse Transcriptase. The relative abundance of *KRAS*, *NRAS* and *HRAS*, gene transcripts was analyzed by using SsoAdvanced Universal SYBR Green Supermix on the CFX-connect real-time PCR instrument (Bio-Rad) and Bio-Rad CFX Manager Software. Specific amplicons were detected for *KRAS* (both K-Ras4A and K-Ras4B splice variants), *NRAS, HRAS*, and *GAPDH*. Forward and reverse primer sequences *KRAS* and *NRAS* amplicons were described previously (Duggan *et al*, 2019; Tsai *et al*., 2015). Primers for amplification of *HRAS* and *GAPDH* were designed using the online tool ‘OligoPerfect Primer Designer’. The mRNA sequences of mouse *HRAS* (NM_008284.3) and *GAPDH* (NM_008084.4) were used as templates for primer design. The relative mRNA expression level was calculated using the 2^-ΔΔCt^ method by normalizing to *GAPDH* expression (Livak & Schmittgen, 2001).

### Data and Statistical analysis

Prism 9 (GraphPad) was used for the preparation of plots, heatmaps, data and statistical analysis. The number n of analyzed cells that was employed for statistical calculations was at least 2000. These originated from N independent biological repeats as indicated in the figure legends. Bar plots show mean ± SD, if not stated otherwise. Statistical analysis of flow cytometry data was performed by employing the Fisher’s Exact test, unless otherwise mentioned in the legends. Immunoblotting data were compared using the unpaired t-test. Half-lives were compared using the Mann-Whitney test.

A p-value < 0.05 is considered statistically significant, and the statistical significance levels are annotated as follows: * p < 0.05; ** p < 0.01; *** p < 0.001; **** p < 0.0001.

## Results

### Benchmarking of flow cytometry-based analysis of C2C12 differentiation

Differentiation stage-specific markers of mouse muscle C2C12 cells correlate with those identified during muscle development *in vivo* (***Figure 1A***) (Yin *et al*., 2013). Both the MAPK-and PI3K-pathways downstream of Ras are characteristically regulated during differentiation (Bennett & Tonks, 1997; Wakioka *et al*., 2001; Xu & Wu, 2000). The C2C12 model therefore offers the opportunity to study the impact of Ras-pathway disease variants and targeted drug treatments on cellular differentiation.

**Figure 1.**
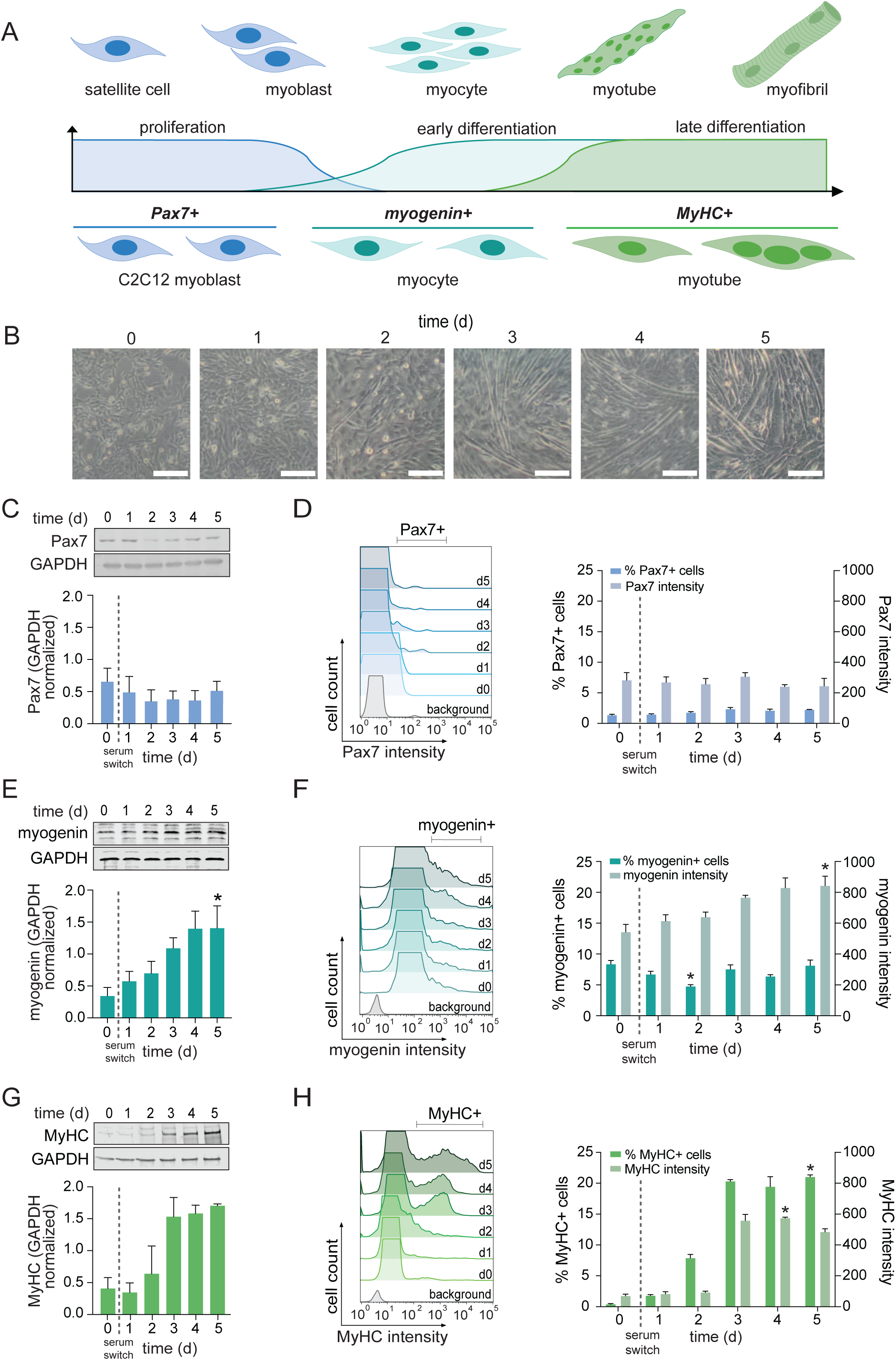
Improved characterization of C2C12 cell differentiation by flow cytometry. (**A**) Differentiation of skeletal muscle *in vivo* (top) and of C2C12 cells (bottom) are similar based on marker expression. (**B**) C2C12 cells were differentiated by switching to low serum medium for indicated times. Brightfield images were obtained at 10 ξ magnification. Scale bars, 200 µm. (**C-H**) Representative immunoblots and quantification of Pax7 (C), myogenin (E) and MyHC (G) expression in C2C12 cells; N = 3, passage number 6. Representative flow cytometry histograms of marker+ cells at indicated times after serum switch (left). Quantification of marker+ fraction and geometric means of intensities (right). Analyzed markers were Pax7 (D), myogenin (F) and MyHC (H); N = 3, passage number 6. Statistical significance was calculated using the Friedman’s test and Dunn’s post hoc test in comparison to day 0. The switch to low serum medium for differentiation is here and subsequently indicated with a vertical dashed line.

We first benchmarked this assay against conventional phenotypic and immunoblotting-based differentiation analysis. C2C12 cell myoblasts assume a roundish-rhomboid morphology when cultured in high serum (day 0) (***Figure 1B***). Medium switching to low serum induces differentiation (day 1), which alters the morphology of cells to become increasingly more spindle shaped. From day 2 to day 3 after serum switching the number of spindle shaped cells visibly increased, and further elongation with subsequent fusion resulted in a robust myotube assembly at day 5 (***Figure 1B***). While determining the fraction of nuclei incorporated into myotubes can measure the overall progression of differentiation as the fusion index, this and related methods are quite tedious and allow only for the analysis of small cell numbers (Velica & Bunce, 2011).

In addition, muscle progenitor and differentiation markers are commonly analyzed using immunoblotting, which however captures only the averaged response across the heterogenous, differentiating cell pool. This became obvious when we compared immunoblotting-and flow cytometry-derived results of various muscle differentiation markers during the 5-day differentiation period. Immunoblotting revealed a nearly constant Pax7 protein expression (***Figure 1C***), which were matched by an almost constant fraction of Pax7 positive (Pax7+) cells that exhibited a constant mean Pax7 intensity in the flow cytometry-based analysis (***Figure 1D***). By contrast, analysis of the early differentiation marker myogenin by immunoblotting suggested a linear increase of the muscle-specific transcription factor during differentiation (***Figure 1E***). This increase is however not due to an increased number of myogenin positive cells, but an increased mean expression level of the transcription factor in the population, as revealed by the flow cytometry data (***Figure 1F***). Both immunoblotting (***Figure 1G***) and flow cytometric analysis (***Figure 1H***) confirmed that the expression of the late differentiation marker myosin heavy chain (MyHC) rapidly increased between days 2 and 3. In this case both the fraction of MyHC+ cells and the mean expression level in the population increased (***Figure 1H***). The flow cytometry-based analysis of C2C12 cell differentiation therefore resolves population level differences in the expression of differentiation markers that may remain hidden in other common types of differentiation analyses. We therefore recently established a protocol to measure C2C12 cell differentiation using the flow cytometric quantification of the MyHC+ fraction, which we furthermore automated by our custom R-script software FlowFate (Parisi *et al*., 2023).

The sensitivity of this assay furthermore enabled us to detect a decline in the differentiation potential with an increase in passage number. Cells with the passage number six (***Figure 1G,H***) arrived at ∼20 % MyHC+ cells at day 3, while this fraction declined to ∼15 % in passage eight and slightly further to ∼13 % in passage ten (***Figure1 – figure supplement 1 A-D***). At the same time, the number of differentiated cells at day 0 in high serum increased in the same order, suggesting that a leakage into differentiation occurs due to passaging even in high serum. As this reduced the net change of the fraction of differentiated cells, i.e. the dynamic range of this assay, we aimed at employing cells with passage numbers from six to nine and included internal references wherever possible. Nevertheless, a residual background fluctuation inherent to variations caused by the passage numbers can be observed in our data that were composed from independent biological repeats across a longer time span.

### Differentiated cells arise from the major pool of Pax7-/ MyHC-transit amplifying cells

Our analysis revealed that the muscle progenitor marker Pax7 is expressed by a minor sub-population of C2C12 cells (< 1%) before and after induction of differentiation (***Figure 2A***). It is therefore impossible that this population of myoblasts provides the bulk of differentiated Pax7-/ MyHC+ myotubes. Instead, double labelling revealed that concomitant with the increase of the Pax7-/ MyHC+ differentiated cells, a Pax7-/ MyHC-subpopulation decreased that constitutes the bulk of the C2C12 cell line (***Figure 2A***). This strongly suggests that the MyHC+ differentiated cells arise from the Pax7-/ MyHC-cells.

**Figure 2.**
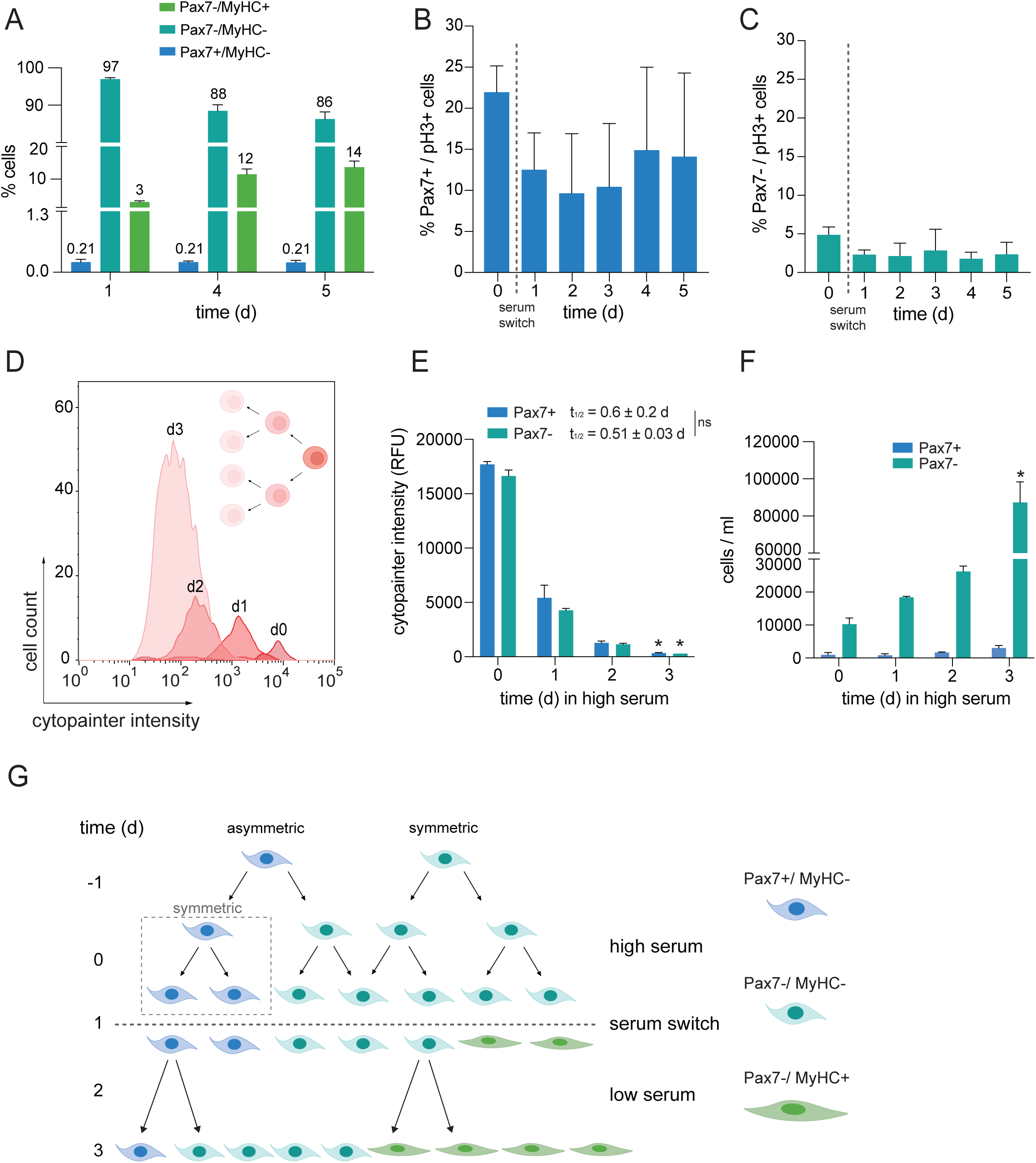
A major pool of Pax7-/ MyHC-transit amplifying cells gives rise to Pax7-/ MyHC+ differentiated cells. (**A**) Flow cytometric analysis of C2C12 cell subpopulations after indicated times of differentiation; N ≥ 2, passage number 8-9. (**B,C**) Quantification of mitotically active (pH3+) Pax7+ (B) and Pax7-(C) subpopulations; N = 4, passage number 7-8. **(D)** Representative histograms of cytopainter dye dilution experiments from C2C12 cells proliferating in high serum medium for indicated times. Inset schematic illustrates dye dilution upon division. **(E)** Quantification of dye dilution experiments as in (D). Geometric means of fluorescence intensities of cytopainter deep red-labelled Pax7+ and Pax7-cells; N = 3, passage number 7-8. Half-life (t_1/2_) of each fluorescence decay was calculated as described in methods. **(F)** Analysis of cell concentrations of Pax7+ and Pax7-populations proliferating in high serum; N = 3, passage number 7-8. **(G)** Revised population model for C2C12 cells. Our data suggest that Pax7+/ MyHC-progenitors divide to some extent symmetrically (boxed), but mostly asymmetrically, to self-renew and expand the number of slowly proliferating Pax7-/ MyHC-transit amplifying cells. This major pool of cells can be triggered to differentiate into Pax7-/ MyHC+ cells in low serum.

To understand how this large pool of Pax7-/ MyHC-cells is maintained under high serum conditions, we analyzed the proliferation of the Pax7+ and Pax7-populations by co-labelling Ser10-phosphorylated histone H3 (pH3), a marker of mitotic and proliferative activity (Goto *et al*, 1999). While ∼22 % of Pax7+ myoblasts were also pH3+ at day 0 in high serum, only ∼5 % of Pax7-cells were pH3+, suggesting that these latter cells have a low mitotic activity. After switching to low serum, the fraction of mitotically active Pax7+ cells dropped to ∼12 % (***Figure 2B***), while the Pax7-population almost stopped dividing (***Figure 2C***).

The higher proliferation rate of Pax7+ cells in high serum was further supported by dye-dilution experiments. The non-toxic cytopainter dye is diluted ∼2-fold during each cell division, which was reflected by the decrease in the geometric means of fluorescence intensities in an exponentially growing cell population (***Figure 2D,E***). When cultured under high serum, the cytopainter labelling decayed at a comparable rate in both Pax7+ and Pax7-subpopulations (***Figure 2E***). At the same time, the number of Pax7-cells increased exponentially during the 3-day culture in high serum, while that of Pax7+ cells only marginally increased (***Figure 2F***). This proliferation pattern is reminiscent of other developmental systems where a minor pool of progenitors asymmetrically divides to generate one progenitor and one transit amplifying cell. The latter then expand exponentially, while the progenitor pool is preserved (Chia *et al*, 2008). This setup is not only critical during development but also for tissue homeostasis and regeneration in the adult (Gomez-Lopez *et al*, 2014; Post & Clevers, 2019).

Our data therefore suggest that under high serum conditions, the Pax7-/ MyHC-transit amplifying cells are replenished via predominantly asymmetric cell divisions of the highly proliferative Pax7+/ MyHC-progenitors, which explains the similar half-life of Pax7+ and Pax7-cell dye dilutions (***Figure 2E,G***). The Pax7-/ MyHC-cells divide slower and likely symmetrically and furthermore continue to be replenished by the asymmetrically dividing progenitors, which explains the exponential expansion of Pax7-cells (***Figure 2F,G***). With terminal differentiation, the Pax7-pool becomes post-mitotic and gradually forms myocytes and myotubes while expressing MyHC (***Figure 2C,G***).

### K-Ras is needed for progenitor maintenance while H-Ras promotes differentiation

It was previously suggested that the three cancer associated Ras genes, *KRAS*, *NRAS* and *HRAS* all promote muscle differentiation via the PI3K-pathway (Lee *et al*, 2010). However, the individual contributions of the Ras genes on the above established three subpopulations of C2C12 cells is unknown. We therefore utilized siRNA-mediated knockdown to specifically downmodulate endogenous *RAS* isoforms and analyzed the effect on the Pax+/ MyHC-progenitors, Pax7-/ MyHC-transit amplifying cells and Pax7-/ MyHC+ terminally differentiated cells during differentiation.

Ras isoform specificity of knockdowns was validated by both immunoblotting (***Figure 3- figure supplement 1***) and quantitative RT-PCR (***Figure 3-figure supplement 2***). This analysis surprisingly revealed that knockdown of *NRAS* significantly increased K-Ras and H-Ras protein expression levels on day 1 (***Figure 3-figure supplement 1F***), which was for K-Ras also reflected on the mRNA-level (***Figure 3-figure supplement 2D***). Hence, this effect is likely not an unspecific siRNA activity, but an endogenous feedback mechanism. In addition, *NRAS* knockdown also downmodulated H-Ras mRNA on day 4 (***Figure 3-figure supplement 2D***). Otherwise, all knockdowns remained isoform specific with an average knockdown efficiency of ≥ 35 % on the protein level during the 5-day differentiation period (***Figure 3-figure supplement 1C,E,G***).

**Figure 3.**
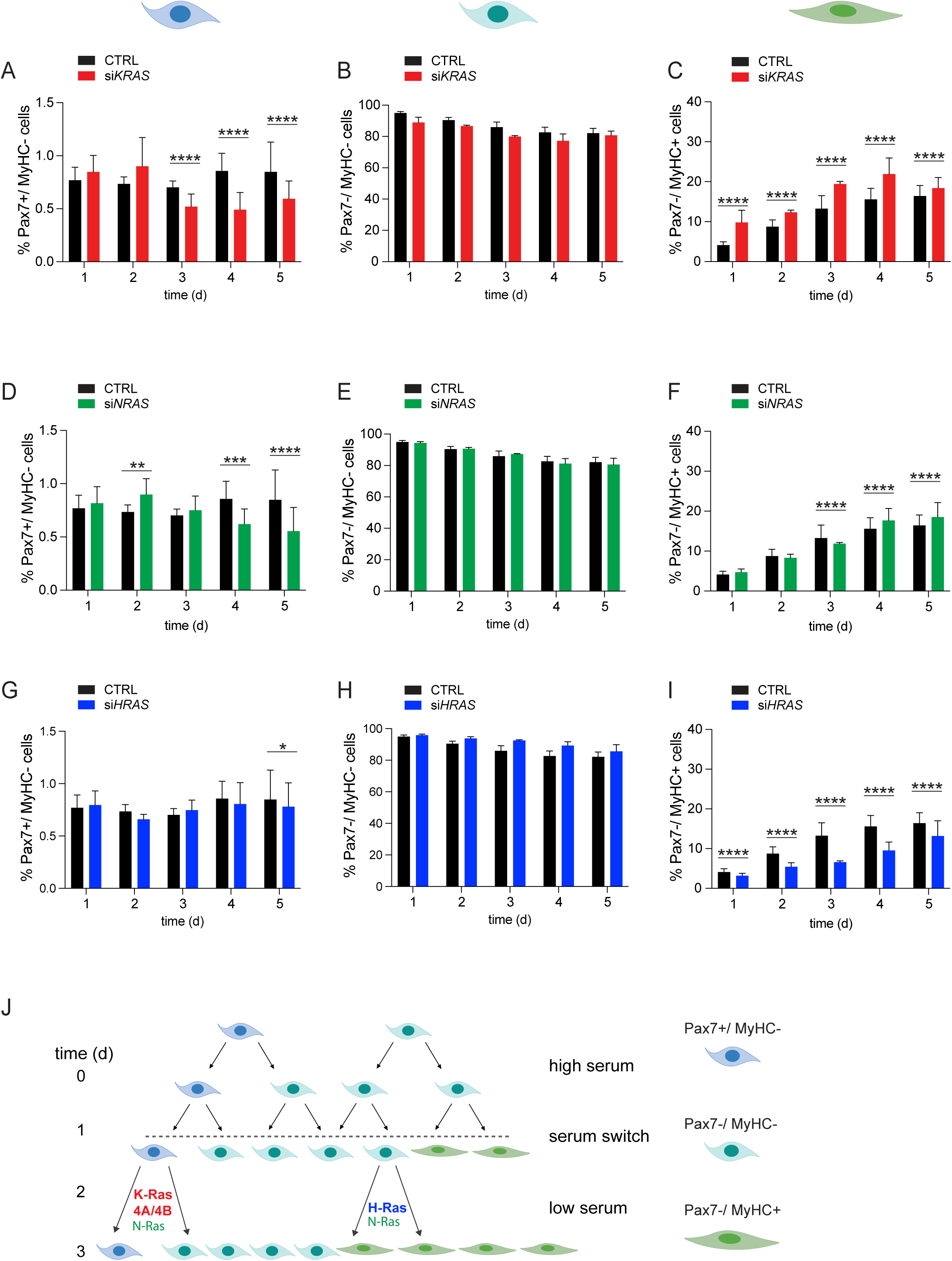
K-Ras4A/4B block whereas H-Ras promotes C2C12 differentiation. (**A**-**I**) Flow cytometric quantification of double-labelled C2C12 cells at indicated times of differentiation. At day 0 cells were treated with si*KRAS* to knock down both isoforms K-Ras4A/4B or control siRNA (CTRL) (A-C). Analogous treatments were done using si*NRAS* (D-F) or si*HRAS* (G-I). Then the fractions of Pax7+/ MyHC-progenitors (A,D,G), Pax7-/ MyHC-transit amplifying cells (B,E,H) and Pax7-/ MyHC+ differentiated cells (C,F,I) were determined by flow cytometric analysis. Ideograms of cells at the top indicate the analyzed cell population in the column of panels; N ≥ 4, passage number 6-7. (**J**) Update of our population model for C2C12 cell differentiation with inferred participation of Ras isoforms in cell divisions.

Specific knockdown of *KRAS* (i.e., of both K-Ras4A and K-Ras4B proteins) led to a significant reduction of progenitors that is more clearly visible from day 3 onwards (***Figure 3A***). Consequently, a slight but not significant drop in the fraction of transit amplifying cells was noticeable (***Figure 3B***). Importantly, the population of differentiated cells was significantly increased upon *KRAS* knockdown (***Figure 3C***). A similar, albeit attenuated effect notably in the population of differentiated cells was observed with the *NRAS* knockdown (***Figure 3D-F***). It is plausible to assume that the smaller effect is due to the *NRAS* knockdown induced upregulation of K-Ras and H-Ras. A markedly different outcome was observed upon knockdown of *HRAS*, which did not markedly alter the progenitor fraction (***Figure 3G***) but slightly increased the fraction of transit amplifying cells (***Figure 3H***), which then resulted in a significantly decreased fraction of differentiated cells (***Figure 3I***).

These population changes are accompanied by a drop in MAPK-and mTORC2-signalling, potentially in an isoform specific manner (***Figure 3 – figure supplement 3***). We observed a decrease of relative Thr202/Tyr204-phosphorylated ERK 1/2 (pERK) levels upon *KRAS* or *HRAS* knockdown, but essentially no change in the case of *NRAS* knockdown (***Figure 3 – figure supplement 3A-D***). Hence, MAPK-signaling is relevant for both progenitor maintenance by K-Ras4A/B (***Figure 3A***) and expansion of differentiated cells by H-Ras (***Figure 3I***). By contrast, downregulation of any *RAS* isoform strongly reduced relative Ser473- phosphorylated Akt1 (pAkt) downstream of Ras and mTORC2 (Kovalski *et al*., 2019), with the strongest effect for *HRAS* knockdown (***Figure 3 – figure supplement 3E-H***).

Taken together, we postulate distinct roles for the three Ras isoforms in regulating C2C12 differentiation (***Figure 3J***). K-Ras4A/B proteins are important to maintain the Pax7+ progenitor pool and prevent differentiation of the transit amplifying cells (***Figure 3A-C***). N-Ras may have a similar and therefore partially redundant role, which may however be obscured given that its downregulation unexpectedly upregulates K-Ras and to a lesser extent H-Ras (***Figure 3D-F***). This may correspond to a fail-safe mechanism, which makes sense in the context that *NRAS* is the evolutionary more recent Ras gene (Garcia-Espana & Philips, 2023).

### Oncogenic K-RasG12V blocks differentiation of transit amplifying cells

It is well established that overactive MAPK-activity, such as associated with disease variants of pathway genes, blocks C2C12 cell differentiation (Konieczny *et al*, 1989; Wakioka *et al*., 2001). However, it is unknown, in which subpopulation these defects manifest and whether proliferation is increased, as typically assumed for oncogenic Ras-transformed cells.

Both oncogenic mutations as well as overexpression of Ras proteins is found in many cancers. Our flow cytometry-based differentiation assay offers the opportunity to exactly quantitate the expression level-dependent effect of both insults on differentiation, by gating for distinct transient expression levels of mEGFP-tagged wild-type or oncogenic K-Ras4B (hereafter K-Ras). Typically, 2000 – 3000 transfected cells were analyzed per condition, a number that remained constant until day 2 of differentiation, when expression started to drop (***Figure 4- figure supplement 1A,B***). This transient perturbation of differentiation is necessary, as in stable transfectants the homeostasis of the mixed C2C12 cell pool would be permanently disrupted. For both constructs, most cells expressed in an expression window, which we defined as up to 10-fold above auto-fluorescence background of untransfected cells (***Figure 4-figure supplement 1C,D***). A high-expression window essentially comprised all cells with expression levels beyond the former threshold.

**Figure 4.**
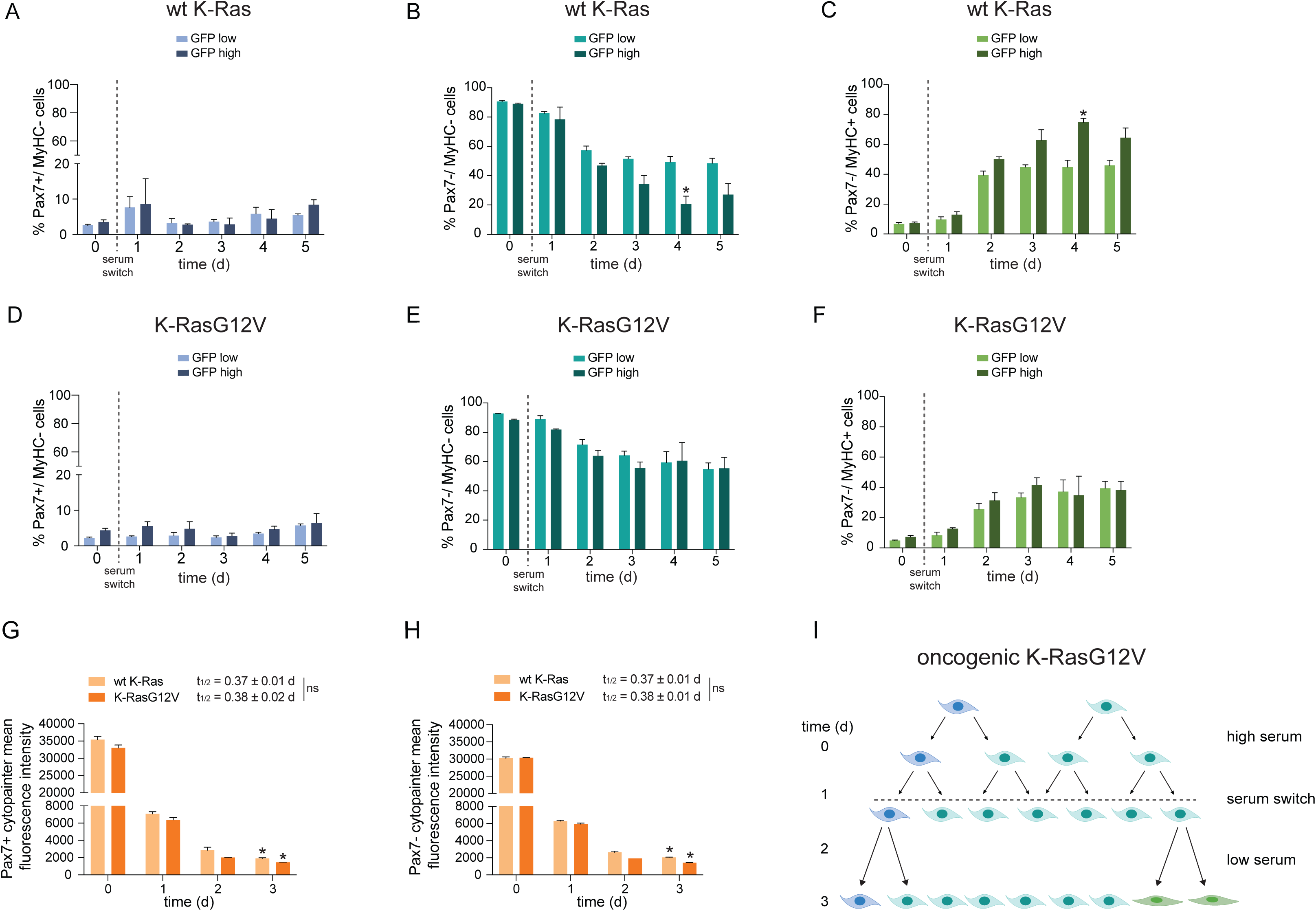
Oncogenic K-Ras4B blocks differentiation of transit amplifying cells. (**A-C**) Flow cytometric analysis of temporal evolution of Pax7+/ MyHC-progenitors (A), Pax7-/ MyHC-transit amplifying cells (B) and Pax7-/ MyHC+ differentiated cells (C) after transfection of C2C12 cells with mEGFP tagged wt K-Ras on day 0. Cells in the GFP low and GFP high expression windows were analyzed; N = 3, passage number 6. (**D-F**) Flow cytometric analysis of temporal evolution of Pax7+/ MyHC-progenitors (D), Pax7-/ MyHC-transit amplifying cells (E) and Pax7-/ MyHC+ differentiated cells (F) after transfection of C2C12 cells with mEGFP tagged oncogenic K-RasG12V on day 0. Cells in the GFP low and GFP high expression windows were analyzed; N = 3, passage number 6. (**G,H**) Geometric means of fluorescence intensities of cytopainter orange-labelled Pax7+ (G) and Pax7-(H) of mEGFP-wt K-Ras or mEGFP-K-RasG12V expressing C2C12 cells; N = 3, passage number 6. Half-life (t_1/2_) of each fluorescence decay was calculated as described in methods. (**I**) Population model for C2C12 cells transformed with oncogenic K-RasG12V, which blocks differentiation at the level of Pax7-/ MyHC-transit amplifying cells.

In C2C12 cells expressing wt K-Ras in the low window, the fraction of Pax7+/ MyHC-progenitors remained essentially constant (***Figure 4A***), and at a comparable level to that of untransfected C2C12 cells (***Figure 1D***). As observed for untransfected C2C12 cells (***Figure 2A***), a decrease in the pool of transit amplifying Pax7-/ MyHC-cells (***Figure 4B***) was matched by an increase in the number of differentiated Pax7-/ MyHC+ cells (***Figure 4C***). When comparing batch and passage number matched cells it can be inferred that the GFP low window corresponds to more physiological expression conditions. In cells expressing wt K-Ras in the high window, the fraction of progenitor cells appeared unaltered (***Figure 4A***). High wt K-Ras expression then resulted in a decreased fraction of transit amplifying cells as compared to the low-expression window (***Figure 4B***), which was matched by an increased fraction of differentiated cells that again increased over time (***Figure 4C***).

The fraction of progenitors appeared likewise unaltered in the GFP low and high windows of cells expressing K-RasG12V (***Figure 4D***). However, the fraction of transformed transit amplifying cells clearly showed a smaller decrease as compared to their wt K-Ras counterparts (***Figure 4E*)**, irrespective of the expression level. This was matched by a reduced increase of differentiated cells (***Figure 4F***).

It is generally assumed that oncogenic Ras mutants increase proliferation, which could impact on the fractions of cells that are analyzed here. We therefore again performed dye-dilution experiments to determine the doubling times in the Pax7+ and Pax7-populations of cells transfected with mEGFP-tagged wt K-Ras and K-RasG12V. Importantly, no significant differences were observed between the cytopainter dye dilution decay half-life values of wt K-Ras and K-RasG12V expressing cells in both the Pax7+ population (***Figure 4G***) and the Pax7-population (***Figure 4H***). K-RasG12V therefore does not stimulate proliferation more than the wt counterpart. Instead, K-RasG12V inhibits differentiation of the transit amplifying cells (***Figure 4I***).

### Oncogenic and RASopathy-associated K-Ras mutants vary in their abilities to block differentiation

We next analyzed the impact of various mutant Ras-alleles on C2C12 cell differentiation. Again, we transiently transfected cells with GFP-variant tagged Ras-constructs and examined them by flow-cytometry in the GFP low window. We focused our analysis on day 3 of differentiation, as it is the earliest time point where differentiation measured by the fraction of MyHC+ cells becomes significantly different (p < 0.0001) between wt and oncogenic K-Ras in the GFP low window (***Figure 4C, F***). In addition to the most frequent K-Ras mutations (Hobbs *et al*, 2016), we also included N-RasG12V and H-RasG12V and their altered biochemical properties for comparison (***Table S1***) (Hunter *et al*, 2015; Moore *et al*., 2020; Rabara *et al*, 2019).

While the Pax7+/ MyHC-progenitor pool appeared slightly increased by most oncogenic Ras mutants (***Figure 5A***), their ability to block differentiation was most distinct (***Figure 5B***). Interestingly, N-RasG12V and H-RasG12V, like K-RasG12D, left the progenitor fraction unaltered (***Figure 5B***), while significantly blocking differentiation similar to all other oncogenic K-Ras mutants (***Figure 5B***).

**Figure 5.**
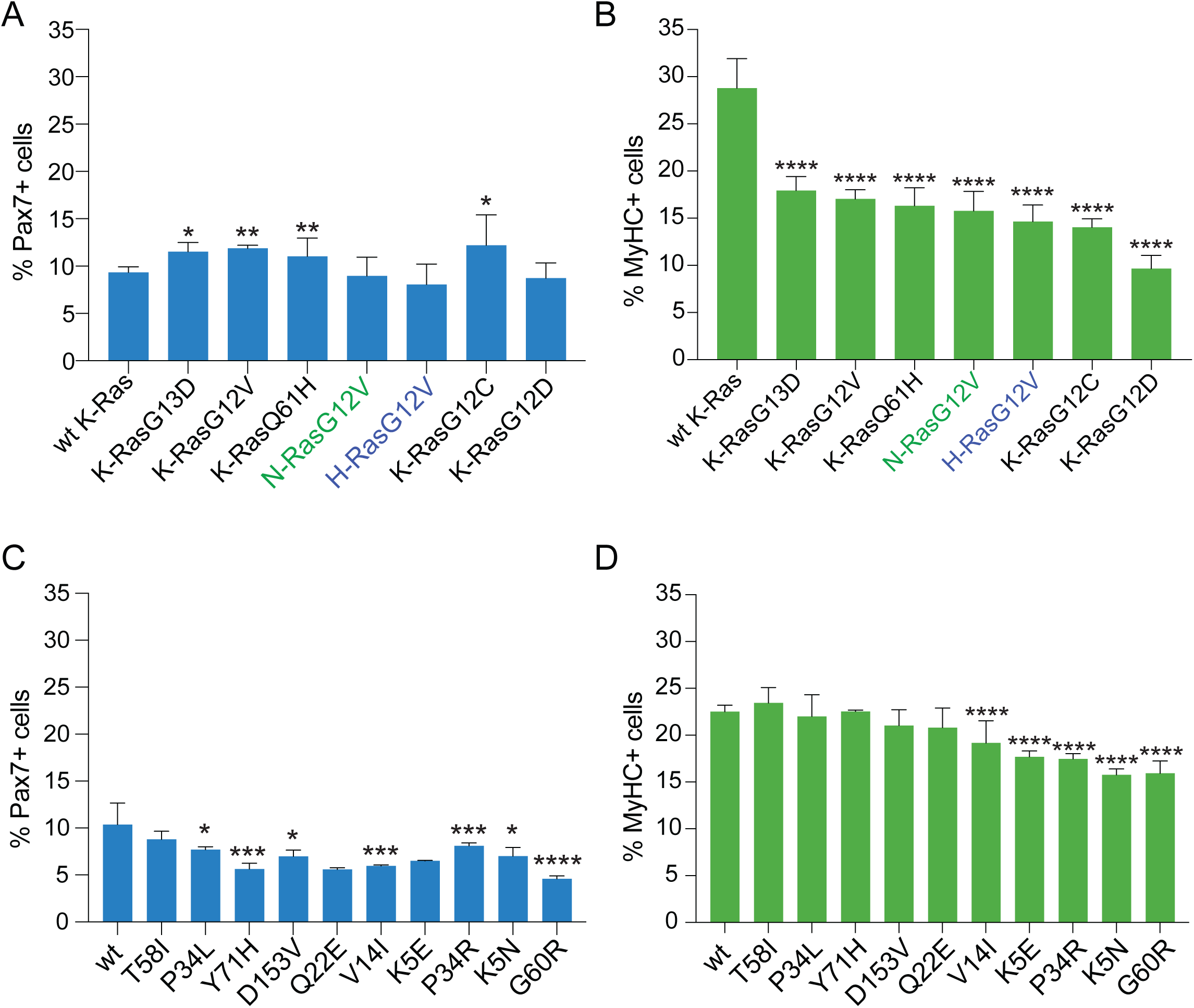
All oncogenic Ras mutants block differentiation, while RASopathy associated K-Ras mutants affect differentiation more broadly. (**A,B**) EGFP-variant tagged oncogenic Ras constructs were transfected into C2C12 cells on day 0 and subsequently on day 3 of differentiation Pax7+ cell fractions (A) and MyHC+ cell fractions (B) were analyzed by flow cytometry; N = 3, passage number 7. (**C,D**) EYFP-tagged K-Ras RASopathy mutants were transfected into C2C12 cells on day 0 and subsequently on day 3 of differentiation Pax7+ cell fractions (A) and MyHC+ cell fractions (B) were analyzed by flow cytometry; N ≥ 2, passage number 9. In all plots, at least 3000 cells in the GFP low expression windows were analyzed.

Aberrant differentiation is also observed in developmental diseases called RASopathies, where Ras-pathway genes are mutated in the germline. Thus, every cell in the body would essentially experience malfunctioning Ras that could broadly impact on development. Consistently, all RASopathies are characterized by multi-organ abnormalities, including of the musculoskeletal system (Stevenson *et al*, 2012). With a few exceptions, *Ras* mutations that are found in RASopathies are different from the ones seen in cancer (Bustelo *et al*, 2018; Castel *et al*., 2020). RASopathy mutants typically display multiple biochemical abnormalities but increase Ras-MAPK signaling less than oncogenic mutants (***Table S2***) (Cirstea *et al*., 2013; Gremer *et al*., 2011).

While the progenitor fractions of K-Ras RASopathy mutant transformed cells were mostly decreased (***Figure 5C***), we saw in general unaltered differentiation or a weaker block of differentiation with RASopathy-derived mutants as compared to oncogenic variants (***Figure 5D***). The most obvious exception was K-Ras with the mutation G60R, which inhibited differentiation similar to the weakest oncogenic mutant K-RasG13D. Five other RASopathy mutants also significantly blocked differentiation and three of these were described to be less NF1-GAP sensitive, a defect that is characteristic for all oncogenic mutants (***Table S1***) (Gremer *et al*., 2011; Schubbert *et al*, 2006). This points to a high significance of the NF1-GAP to facilitate terminal differentiation of transit amplifying cells and implicitly to a disruption of such differentiation steps in general during Ras-driven oncogenesis.

### K-RasG12C inhibitor profiling reveals their distinct ability to restore differentiation and induce toxic cell death

The past few years have seen the arrival of the first direct K-RasG12C inhibitors in the clinic (Punekar *et al*, 2022; Steffen *et al*., 2023). Classical cell-based assays profile these inhibitors based on their anti-proliferative and cell-killing activity. Here, we have the unique opportunity to quantify to what extent these inhibitors can restore inhibited differentiation that was induced by oncogenic K-Ras alleles.

First, we established that the employed DMSO concentrations below 0.1 % that carried over from compound stocks do not impact on differentiation or have toxic effects (***Figure 6-figure supplement 1A,B***).

**Figure 6.**
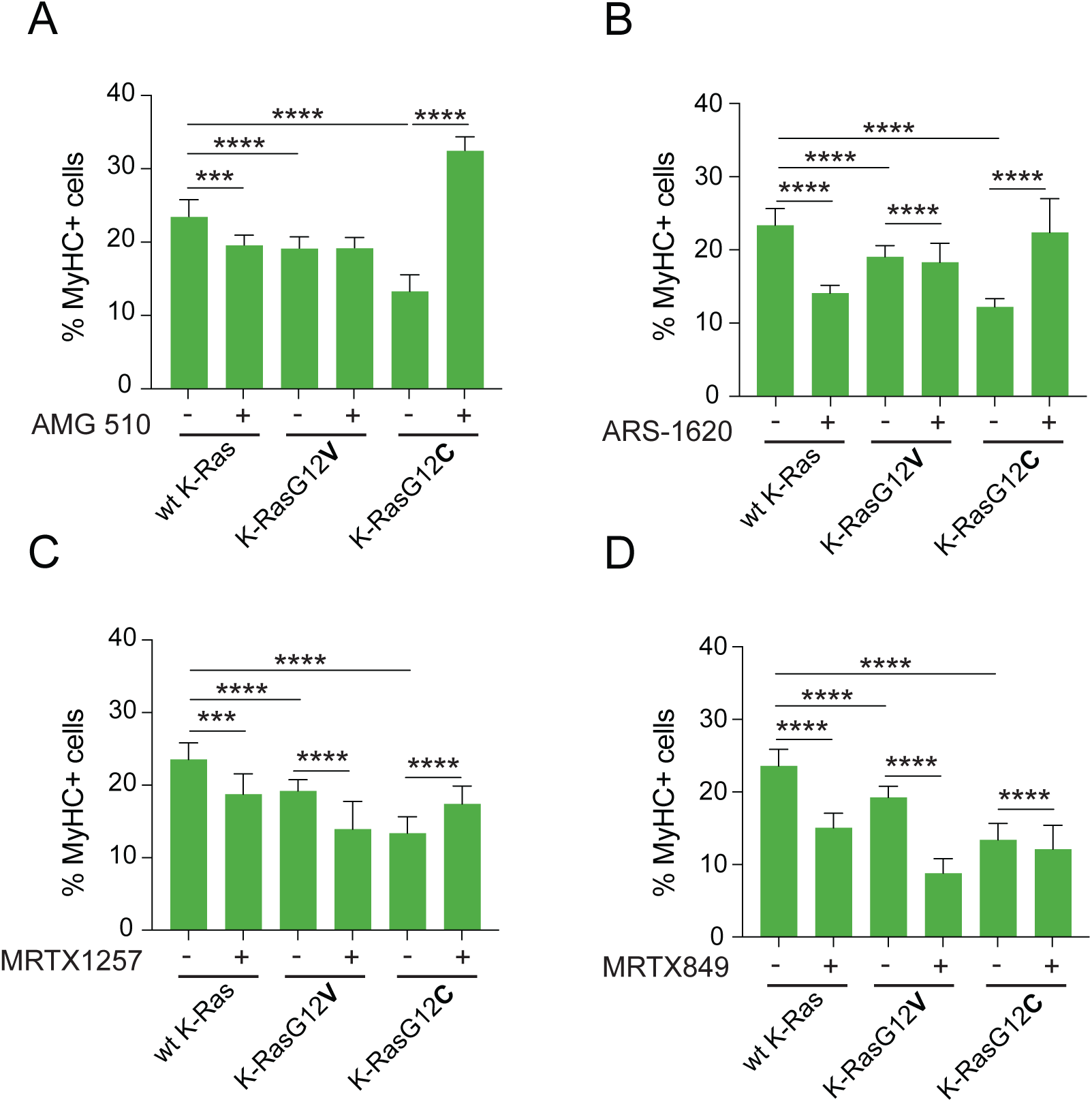
K-RasG12C inhibitor profiling reveals their distinct ability to restore differentiation. (**A-D**) Analysis of MyHC+ C2C12 cell fractions expressing GFP-variant tagged wt K-Ras, K-RasG12V and K-RasG12C on day 3 of differentiation. From day 1 onwards 3 μM of AMG 510 (A), ARS-1620 (B), MRTX1257 (C) or MRTX849 (D) or vehicle (DMSO 0.1 % v/v) were added to cells; N = 4. In all plots, only cells from passage 9 in the GFP low expression windows were analyzed.

We then tested the two approved drugs sotorasib (AMG 510) and adagrasib (MRTX849) (Canon *et al*., 2019; Fell *et al*., 2020), as well as ARS-1620 the founder of current G12C-inhibitors (Janes *et al*., 2018), and MRTX1257, a close analogue of MRTX849 (Fell *et al*., 2020).

As just described, K-RasG12C decreased the fraction of MyHC+ cells more than K-RasG12V on day 3 (***Figure 6A***). AMG 510 treatment at 3 μM did not have a restorative effect on differentiation of cells expressing wt K-Ras or K-RasG12V, however, it significantly increased the fraction of differentiated cells with K-RasG12C even beyond that of the wt-control (***Figure 6A***). This was an interesting observation, as it may indicate a dominant negative action of AMG 510-bound K-RasG12C since this phenotype resembles that of the K-RasG12V-S17N with the S17N dominant negative mutation (***Figure 6-figure supplement 1C,D***). This distinct ability of AMG 510 has not been reported previously. As expected, ARS-1620 could fully restore differentiation of K-RasG12C transformed C2C12 cells (***Figure 6B***), while having no effect on the number of intact cells as also observed with AMG 510 (***Figure 6-figure supplement 1E,F***).

By contrast, neither MRTX1257 nor MRTX849 could fully restore differentiation of K-RasG12C transformed C2C12 cells (***Figure 6C,D***). Instead, we observed a reduction in the MyHC+ fraction in the drug treated K-RasG12V expressing cells (***Figure 6C,D***). This was due to significant general toxicity of these compounds, which led to a significant drop of intact cells even for wt K-Ras expressing cells (***Figure 6-figure supplement 1G,H***).

We more closely examined this K-RasG12C-inhibitor toxicity using 7-AAD-labelling, which indicates late apoptosis and necrosis in cells (Zembruski *et al*, 2012). This assay confirmed that under otherwise the same conditions the general toxicity increased in the order AMG 510 ≲ ARS-1620 < MRTX1257 ≲ MRTX 849 in adherent and detached K-RasG12C transformed C2C12 cells (***Figure 6-figure supplement 1I,J***).

This analysis demonstrates that both AMG 510 and ARS-1620 are effective and specific K-RasG12C inhibitors that can restore differentiation with no or little non-specific toxicity. On the other hand, both MRTX1257 and MRTX 849 display a broader toxicity, which undermines their differentiation restoring ability.

### Profiling of clinical and pre-clinical Ras inhibitors for their ability to restore differentiation

In extension of this analysis, we next assessed the oncogene-and allele-specific effect of targeted drugs on differentiation. Altogether we tested eight approved or clinically evaluated Ras-pathway inhibitors at 1 μM (except for trametinib at 0.1 μM and AMG 510 at 3 μM), which target Ras trafficking (tipifarnib and cysmethynil), upstream activation (gefitinib, BI-3406), directly K-Ras (AMG 510, MRTX1133) and the major effector pathways downstream of Ras that are associated with cancer (trametinib, rapamycin).

Trafficking inhibitor tipifarnib, which inhibits farnesyltransferase, had a surprisingly broad effect on almost all K-Ras, N-Ras and H-Ras mutants to restore differentiation (***Figure 7A***). By contrast, cysmethynil was essentially inefficacious (***Figure 7A***), in agreement with its clinical performance (Lau *et al*, 2014).

**Figure 7.**
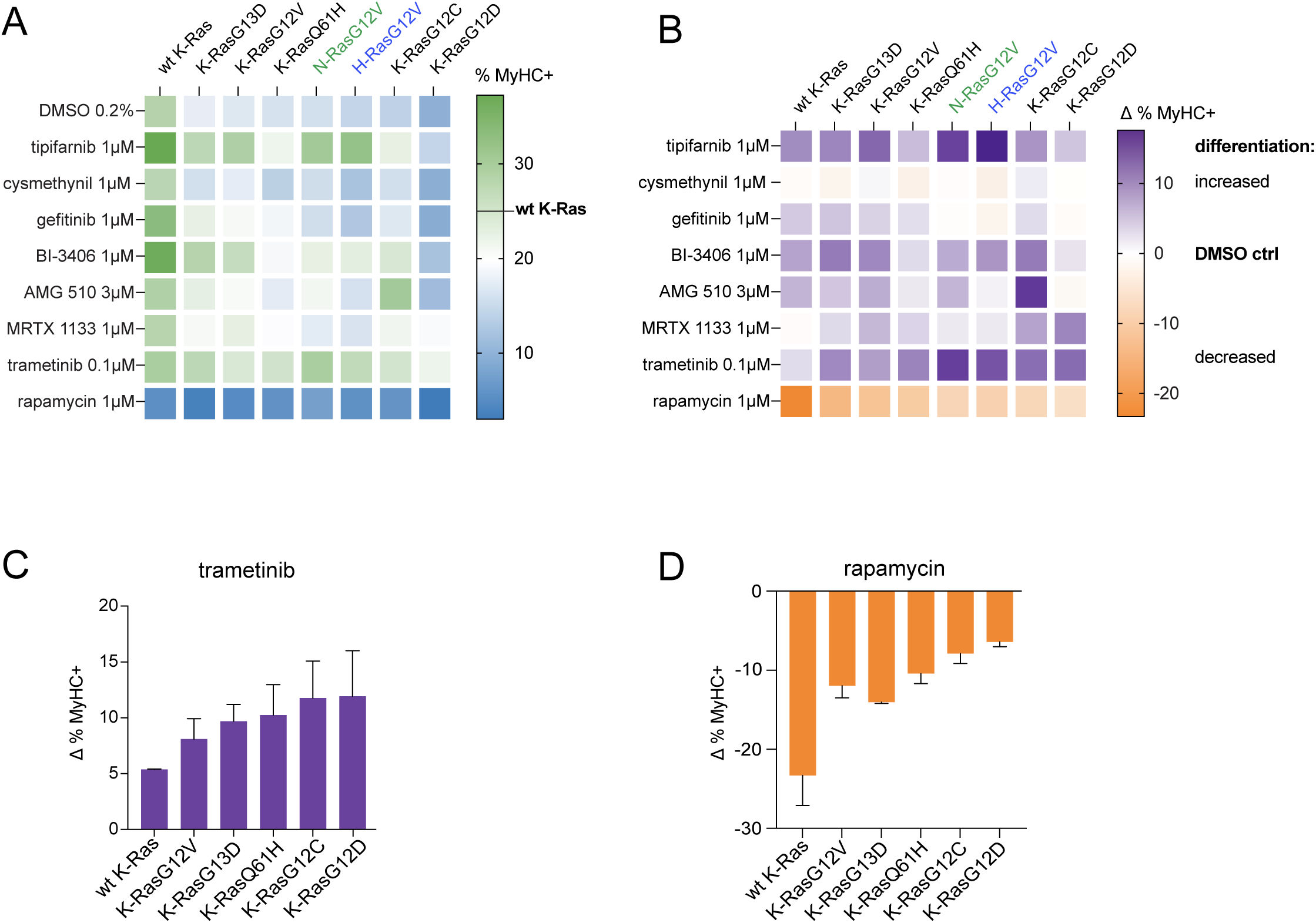
Profiling of clinically tested Ras inhibitors for their ability to restore differentiation. (**A**) Heatmap showing % MyHC+ C2C12 cells, i.e. the absolute rescue effect, on day 3 of differentiation for each indicated condition. EGFP-variant tagged Ras-constructs were transfected on day 0; N = 3, passage number 7. The scale to the right shows the wt K-Ras differentiation level as a reference. (**B**) Net rescue effect of drugs. Data from (A) were used by subtracting values from vehicle control (DMSO 0.2 % v/v) treated samples in the top row from those treated with drugs. The scale to the right shows the DMSO-control level as a reference. (**C**) Extract of data from (B). MAPK-pathway dependence of K-Ras variants judged by the trametinib induced net rescue. (**D**) Extract of data from (B). PI3K/ mTORC1-pathway dependence of K-Ras variants judged by the rapamycin induced net rescue. In all plots, only cells in the GFP low expression windows were analyzed.

The expected H-Ras specificity of tipifarnib treatment becomes more clearly visible, if we consider the net rescue effect of differentiation, i.e. the difference between the MyHC+ fraction with drug treatment and the DMSO-control (***Figure 7B***). The second most sensitive allele was N-RasG12V, interestingly followed by K-RasG12V, while all other oncogenic K-Ras variants were less sensitive. This was surprising, given that K-and N-Ras can undergo its function-restoring alternative prenylation if farnesyltransferase is inhibited (Whyte *et al*, 1997).

Upstream inhibitors gefitinib, which blocks EGFR tyrosine kinase activity, only had a modest effect to restore differentiation of some K-Ras alleles (***Figure 7A,B***), which essentially followed the order of the differentiation blocking activity of these K-Ras mutants (***Figure 5B***). The SOS1 inhibitor BI-3406 performed better, leading to a substantial restoration, except for K-RasQ61H, which is known to have an elevated nucleotide exchange activity that allows its activation independent from SOS1 (Gebregiworgis *et al*, 2021; Hofmann *et al*, 2021). By contrast, this compound was more efficient against K-RasG13D with a much-increased nucleotide exchange activity.

Next, AMG 510 showed the expected selectivity for K-RasG12C, with a minor effect on other alleles (K-RasG13D, K-RasG12V and N-RasG12V) (***Figure 7A,B***). Allele selectivity, albeit not as clear as with AMG 510, was also seen with the low nanomolar non-covalent K-RasG12D-selective inhibitor MRTX1133 (Wang *et al*., 2022). Importantly, MRTX1133 restored differentiation to the highest levels observed so far for K-RasG12D but did appear to also mildly and non-specifically restore the differentiated fraction in other K-Ras alleles (***Figure 7A,B***). These apparent non-specific effects of the two allele specific inhibitors may arise from non-covalent interactions with the switch II pocket of all K-Ras alleles. For compounds with the MRTX849 scaffold such non-specificity was explicitly demonstrated earlier (Vasta *et al*, 2022).

None of the former compounds though exerted an as strong pan-Ras effect as trametinib, which broadly restored differentiation across all alleles and isoforms (***Figure 7A,B***). Only, drugs that have a known allele-or isoform-selectivity (tipifarnib for H-RasG12V, AMG 510 for K-RasG12C and MRTX1133 for K-RasG12D) matched or surpassed the effect of trametinib. This correlated with the clinical success of tipifarnib against H-RasG12V driven tumors and of AMG 510 against K-RasG12C driven tumors (Ho *et al*, 2021; Skoulidis *et al*, 2021). Finally, rapamycin broadly blocked differentiation, consistent with the importance of PI3K/ mTORC1- signaling for myogenesis and terminal differentiation of C2C12 cells (Lee *et al*., 2010; Xu & Wu, 2000).

We therefore, recapitulate closely the *in vivo* effects of targeted pharmacological agents. The broad effect of trametinib moreover highlights the central role of MAPK-activity to block differentiation. This becomes even clearer, when the net restoration of all oncogenic K-Ras alleles by trametinib is analyzed (***Figure 7C***). One recovers a rank-order that anti-correlates with that of rapamycin (***Figure 7D***) and the impact of these alleles on differentiation (***Figure 5B***). This suggests that those K-Ras mutants that have a stronger dependence on MAPK-signaling, as expressed by their relative trametinib sensitivity (***Figure 7C***), and lesser dependence on the PI3K/ mTORC1-signaling (***Figure 7D***), more profoundly block differentiation (***Figure 5B***).

## Discussion

The impact of Ras-MAPK-pathway disease mutants on cell transformation and the efficacy of novel Ras drug candidates are typically assessed in a small number of standard assays, such as NIH3T3 transformation and 2D/3D cancer cell proliferation assays, complemented by immunoblotting for markers of Ras-MAPK-pathway activity (Esposito *et al*, 2019). The impact on cell differentiation, though a known hallmark of cancer, has so far been vastly neglected (Chaffer & Weinberg, 2015; Hanahan, 2022).

Here we have established the C2C12 cell differentiation model to examine the effect of oncogenic Ras mutants and Ras-pathway drugs on differentiation. Importantly, this model recapitulates typical and fundamental steps of cell differentiation also found *in vivo* (Yin *et al*., 2013). Notably, the NF1 scaffold SPRED1 is induced upon differentiation, concomitant with a drop in MAPK-signaling, which allows to examine this core machinery of Ras-transformation in more detail (Stowe *et al*., 2012; Wakioka *et al*., 2001). This implies that our observations have broad implications not only for locally perturbed differentiation in cancer, but also for RASopathies, which are caused by aberrant Ras signaling throughout development. Our approach with a standard commercial cell line is advantageous as compared to the usage of human embryonic stem cells (hESC), which have also been used to identify compounds that maintain stemness or promote differentiation in more laborious, imaging-based high content screens (Barbaric *et al*, 2010; Desbordes *et al*, 2008; Jee *et al*, 2012; Sherman & Pyle, 2013).

Based on our analysis, we propose a new model on how in the C2C12 cell culture a small number of Pax7+ progenitor myoblasts are maintained, while the majority of cells is primed to differentiate. We suggest that frequent asymmetric divisions maintain the Pax7+/ MyHC-progenitor pool, while generating a transit amplifying Pax7-/ MyHC-pool of cells that exponentially expands within a few symmetric divisions. We furthermore show here that the three cancer associated Ras genes, *KRAS*, *NRAS* and *HRAS* have a distinct, yet partially overlapping involvement in sustaining the proper trajectory from progenitors via here identified transit amplifying cells to differentiated cells. Individual knockdowns generate distinct changes in the three populations that can hardly be explained by the alternative scenario where total Ras levels determine these alterations. Our data suggest a particular relevance of MAPK-signaling for K-Ras4A/B-dependent progenitor maintenance and H-Ras-dependent terminal differentiation. This is consistent with previous reports of MAPK inhibition resulting in increased differentiation (Rommel *et al*, 1999). Our model adds another dimension to currently debated models for the isoform-specific functions of these *Ras* genes, that include differences in plasma membrane organization and effector usage (Mo *et al*, 2018). The generally observed high expression level in particular of K-Ras4B in almost all tissues is consistent with its major function to sustain progenitors and being the evolutionary most ancient Ras isoform (Garcia-Espana & Philips, 2023; Hood *et al*., 2023).

It is well established that oncogenic Ras, constitutively active Raf or MEK1 inhibit myoblast differentiation (Dorman & Johnson, 1999; Olson *et al*, 1987; Weyman & Wolfman, 1998). However, we observed two distinct types of perturbed differentiation by Ras-pathway hyperactivation. The first is associated with overexpression of wt K-Ras that led to a more pronounced drop in the transit amplifying population matched by a significantly increased differentiation. By contrast, all NF1-GAP insensitive oncogenic Ras variants blocks terminal differentiation as compared to wt K-Ras. Moreover, those mutants that show a stronger sensitivity to trametinib and a lower sensitivity to rapamycin, are more potent to block differentiation. This order does not exactly fit with the mutation frequency of the oncogenic K-Ras alleles, but it could reflect their severity as expressed by the overall survival associated with the mutants. And exception is K-RasG13D, which is a biochemical outlier with a high nucleotide exchange rate, high intrinsic GTPase activity and NF1-GAP sensitivity (Hunter *et al*., 2015; Rabara *et al*., 2019).

As already inferred from their biochemical characterization, RASopathy associated K-Ras mutants are not or less able to block differentiation, consistent with these alterations being compatible with organismal development. Instead, it appears that many of them are defective in sustaining the C2C12 cell progenitor fraction. Intriguingly, the most potent RASopathy mutant K-RasG60R, which inhibits differentiation as much as the weaker oncogenic K-Ras allele K-RasG12V, also displays a more prominent NF1-GAP resistance. The milder effects of RASopathy K-Ras mutants G60R and P34R can be explained by their decreased effector engagement, which also exists in a milder manifestation in the V14I mutant. We postulate that both K5-mutants also impact on NF1-GAP activity in the cell, given their ability to significantly block differentiation.

We speculate that for all of these NF1-GAP resistant mutants, differentiation of transit amplifying cells is blocked, as shown for K-RasG12V. This is consistent with the fact that with the induction of differentiation, the potentially K-Ras-selective tumor suppressor complex of SPRED1 with the GAP NF1 can become active only after SPRED1-induction (Siljamaki & Abankwa, 2016; Stowe *et al*., 2012; Wakioka *et al*., 2001). Thus, MAPK-signaling would be sustained in transit amplifying cells, which prevents differentiation. This specifies that the oncogenic insult of hotspot-mutated Ras occurs at a defined point of the differentiation trajectory, an important fact that has not been recognized before. Moreover, this model implies molecular mechanistic and developmental commonalities between cancer and RASopathies, which have been long elusive (Castel *et al*., 2020).

In direct correlation with our muscle cell line observations, RASopathy patients display muscle weakness in particular in Costello syndrome (CS) (Stevenson *et al*., 2012; Stevenson & Yang, 2011). In the heterozygous G12V CS mouse model a decreased muscle mass and strength was found, due to inhibited embryonic myogenesis and myofiber formation (Tidyman *et al*, 2022). This was due to an inhibited differentiation in the embryonic muscles, with a 23 % increase in Pax7 expressing cells and a decrease in MyoD and myogenin expressing cells to 60-70 % of the wt. A less severe skeletal myopathy is observed in the RASopathy cardiofaciocutaneous syndrome mouse model with a BRAF-L597V mutation (Maeda *et al*, 2021). Given the distinct penetrance of muscle phenotypes in RASopathies, one may assume this could be due to distinct mutant allele strengths, as suggested by our data. Alternatively, in muscle cells only certain alleles could become significant, while others would be tissue specifically contained, which is probably a less likely scenario. Importantly, these data corroborate the idea that the muscle phenotype in RASopathies is due to perturbed differentiation of stem/ progenitor cells, as has been observed in other muscle diseases notably in Duchenne muscular dystrophy, where asymmetric cell divisions of satellite cells are likewise not proceeding correctly (Feige *et al*, 2018).

Soft tissue rhabdomyosarcoma (RMS) of the muscle are frequently observed in RASopathies, such as Neurofibromatosis type 1, Noonan syndrome and CS (Skapek *et al*, 2019). RMS is the most common childhood soft-tissue sarcoma with only 30 % survival in the metastatic disease. These tumors emerge from muscle progenitors/ myoblasts that failed to differentiate, albeit the exact cell of origin is still not well characterized (Skapek *et al*., 2019). Strikingly, this is exactly the phenotype we have observed in our data.

Two RMS subtypes are distinguished, the alveolar type in adolescents and the embryonal type in younger patients, which is associated with good prognosis, despite higher mutational burden (Shern *et al*, 2014). The former largely overlaps with the Pax-fusion positive molecular subtype, with neomorphic gain-of-function fusion proteins of Pax3 or Pax7 with FOXO1. In the Pax-fusion negative (embryonal) subtype of RMS, the Ras-pathway is activated by mutations in the pathway, while in the Pax-fusion positive subtype the upregulation of Ras pathway genes is found (Shern *et al*., 2014; Skapek *et al*., 2019). Interestingly, it is one of the few cancer types where the three Ras isoforms are mutated at about equal frequency. Oncogenic Ras prevents differentiation in rhabdomyosarcoma (Yohe *et al*, 2018).

Finally, we profiled the effect of drugs on differentiation in a rapid and unique manner. Our assessment of four K-Ras-G12C inhibitors importantly demonstrates that inhibition of the oncogenic K-Ras also rescues differentiation. Yet, unexpected idiosyncrasies of these compounds were observed. Interestingly, the increase in differentiation that we observed with K-RasG12C bound AMG 510, may suggest a molecular complex that becomes dominant negative, similar to what is seen with S17N-mutation. It is not likely that target-independent, off-target effects are responsible for this observation, as there is no effect with K-RasG12V expressing cells. By contrast, adagrasib (MRTX849) and more so MRTX1257 were less proficient in restoring differentiation, while inducing significantly more non-specific cell death. This is typically not desired, and in this particular case it may be attributed to the inhibition of wt K-Ras (Vasta *et al*., 2022). Their inhibition of other mutant *KRAS* alleles is probably beneficial in the clinical setting, where any antiproliferative activity could be helpful and the activity against other *KRAS* alleles prevents their success if they evolved as a resistance mechanism (Liu *et al*, 2022).

Consistent with observations in the more complex hESC model and zebra fish larvae (Chen *et al*, 2011; Pal *et al*, 2012), we also noted the significant, dose-dependent effect of DMSO at concentrations above 0.2 % on C2C12 cell differentiation and viability. Hence, our assay may be suitable to identify and explain differentiation perturbing, toxic effects of organic solvents or other substances that may correlate with their teratogenic potential.

We furthermore illustrate the enormous potential for Ras-pathway drug profiling on multiple Ras-disease mutants in our 8 × 8 matrix, which revealed a remarkable correlation of the ability of drugs to restore differentiation with their clinical efficacy in Ras disease treatment. Trametinib was very efficacious to restore differentiation of K-Ras mutants that profoundly blocked differentiation. Interestingly, the same order of sensitivity was found for tipifarnib, which may indicate that both drugs, tipifarnib and trametinib could work synergistically to block differentiation. The broad capacity to restore differentiation by MEK-inhibition is paralleled by results obtained with myocyte cultures derived from the CS mouse model. Differentiation of those cells could be restored by the MEK inhibitor PD0325901, while the differentiation of the wt control was further increased. Importantly, this was mirrored by muscle-mass and -diameter increases *in vivo* to control levels (Tidyman *et al*., 2022). Thus, relative muscle strength development in RASopathy patients may also serve as a biomarker for treatment efficacy. We found that rapamycin was one of the few compounds, which prevented C2C12 cell differentiation. Others have previously shown that it inhibits differentiation that is promoted by the PI3K/ mTORC1- pathway (Hatfield *et al*, 2015). While the PI3K/ Akt/ mTORC1/ S6K1-axis is involved in hypertrophic muscle growth (Mounier *et al*, 2011; Yoon, 2017), the PI3K inhibitor GDC0941 led to muscle cell death in a CS mouse model and was deleterious *in vivo* (Tidyman *et al*., 2022). However, in the aging muscle, hyperactive mTORC1 appears to induce muscle damage and loss, hence low dose treatment with rapamycin analogue everolimus (RAD001) salvages this situation *in vivo* (Joseph *et al*, 2019). In the end, this compound analysis allowed us to estimate the utilization of the MAPK-and PI3K-pathway suggesting that oncogenic K-Ras alleles show a distinct dependence on these pathways.

Interestingly, those mutants with a higher MAPK (trametinib)- and lower PI3K/ mTORC1 (rapamycin) -dependence were more potent to block differentiation.

Future developments of such drug-profiling derived models may enable us to predict efficacious drug combination that do not only restore differentiation but in parallel also work in cancer therapy. The fact that we observe striking correlations of our differentiation data with overall Ras mutation strength and drug-responses observed in the clinic, suggests that we are looking here at a highly conserved function of Ras that is deeply engrained into the functioning of nearly every metazoan cell system. In order to devise therapies that can fully salvage the aberrant differentiation induced by Ras-pathway hyperactivation, we need to understand its impact on both cell proliferation and differentiation. The involvement of Ras beyond the G1- phase, where the Ras-MAPK pathway is known to drive S-phase entry such as by stimulating cyclin D expression, is largely unknown. However, cell fate decisions are taken during M-phase, as stem/ progenitor cells symmetrically or asymmetrically divide. We therefore postulate a distinct role of Ras-pathway activity during this fundamental step, which cannot merely be explained by different strengths of the pathway output, but the underlying cell and developmental biology.

## Resource availability

### Lead contact

Further information and requests for resources and reagents should be directed to and will be fulfilled by the lead contact, Daniel Kwaku Abankwa.

### Materials availability

This study did not generate new unique reagents.

### Data and code availability

No standardized datasets were produced within this study.

## Acknowledgements

This work was supported by a grant from the Luxembourg National Research Fund (FNR) grant C19/BM/13673303-PolaRAS2 to D.K.A.

## Author contributions

R.C. and D.K.A participated in the initial conceptualization. R.C. and B.P. designed and carried out flow cytometry experiments and data analysis. C.L collected and evaluated immunoblot and qPCR data for RAS isoform knockdown. F.K acquired immunoblot data for differentiation marker expression and performed 7AAD-cell toxicity assay. N.B.F generated expression constructs via gateway cloning. R.C., B.P. and D.K.A wrote the first draft of the manuscript.

D.K.A. edited the manuscript and supervised the entire study.

## Declaration of interests

The authors declare no competing interests.

## Notes

### Competing Interest Statement

The authors have declared no competing interest.

### Summary of Updates

Introduction: The following sentences along with the corresponding references have been added: Ras can furthermore activate mTORC2, which phosphorylates Akt on Ser473, and: Terminal differentiation is then promoted by mTORC2-Akt activity. Figures/tables and legends: C2C12 differentiation schematic in Fig 2G has been updated. Slopes and numbers atop bar blots have been removed in the updated Figure 3. Figure 4 has been modified to remove the signaling schematic in G and all datasets are now presented in Figure 3- supplement 3. Order of the subsequent figures has thus changed and Figure 5 in the previous version is now Figure 4. Figure 8 (which is Figure 7 in the current version) has been revised to exclude schematics and coloring of heatmaps and bar plots has changed. Data in Figure 6 supplement 1I,J have been revised. Corresponding legends have been updated in lieu of changes to the figure content and figure numbering. Supplementary table S1 has been split into tables S1 which represents biochemical properties of oncogenic K-Ras mutants and table S2, which represents biochemical properties of RASopathy associated K-Ras mutants. Corresponding legends have also been updated. Results: The average knockdown efficiency for data in Figure 3 supplement 1 has been revised from >50% to >35%. The subsection: K-Ras4A/B sustain MAPK-signaling, while H-Ras sustains both MAPK- and PI3K-signalling during differentiation, has been removed and relevant portions have been integrated into the sub-section: K-Ras4A/B are needed for progenitor maintenance while H-Ras promotes differentiation. Methods: The source of pmEGFP-KRas4B-G12V-S17N expression construct has been revised. Flow cytometry section has been updated to include the procedure used to generate histograms. Statistical analysis section has been revised to include unpaired t-test. Cell toxicity analysis with 7-AAD sub-section has been revised corresponding to changes in figure 6 supplement 1.

## References

(2022) The KRAS(G12D) inhibitor MRTX1133 elucidates KRAS-mediated oncogenesis. Nat Med 28: 2017-2018

Abankwa D, Gorfe AA, Inder K, Hancock JF (2010) Ras membrane orientation and nanodomain localization generate isoform diversity. Proc Natl Acad Sci U S A 107: 1130–1135

Ahmadian MR, Stege P, Scheffzek K, Wittinghofer A (1997) Confirmation of the arginine-finger hypothesis for the GAP-stimulated GTP-hydrolysis reaction of Ras. Nat Struct Biol 4: 686–689

Altshuler A, Verbuk M, Bhattacharya S, Abramovich I, Haklai R, Hanna JH, Kloog Y, Gottlieb E, Shalom-Feuerstein R (2018) RAS Regulates the Transition from Naive to Primed Pluripotent Stem Cells. Stem Cell Reports 10: 1088–1101

Ansieau S (2013) EMT in breast cancer stem cell generation. Cancer Letters 338: 63–68

Barbaric I, Gokhale PJ, Jones M, Glen A, Baker D, Andrews PW (2010) Novel regulators of stem cell fates identified by a multivariate phenotype screen of small compounds on human embryonic stem cell colonies. Stem Cell Res 5: 104–119

Barretina J, Caponigro G, Stransky N, Venkatesan K, Margolin AA, Kim S, Wilson CJ, Lehar J, Kryukov GV, Sonkin D et al (2012) The Cancer Cell Line Encyclopedia enables predictive modelling of anticancer drug sensitivity. Nature 483: 603–607

Batlle E, Clevers H (2017) Cancer stem cells revisited. Nat Med 23: 1124–1134

Bennett AM, Tonks NK (1997) Regulation of distinct stages of skeletal muscle differentiation by mitogen-activated protein kinases. Science 278: 1288–1291

Brems H, Pasmant E, Van Minkelen R, Wimmer K, Upadhyaya M, Legius E, Messiaen L (2012) Review and update of SPRED1 mutations causing Legius syndrome. Hum Mutat 33: 1538–1546

Brown DM, Parr T, Brameld JM (2012) Myosin heavy chain mRNA isoforms are expressed in two distinct cohorts during C2C12 myogenesis. J Muscle Res Cell Motil 32: 383–390

Bustelo XR, Crespo P, Fernandez-Pisonero I, Rodriguez-Fdez S (2018) RAS GTPase-dependent pathways in developmental diseases: old guys, new lads, and current challenges. Curr Opin Cell Biol 55: 42–51

Canon J, Rex K, Saiki AY, Mohr C, Cooke K, Bagal D, Gaida K, Holt T, Knutson CG, Koppada N et al (2019) The clinical KRAS(G12C) inhibitor AMG 510 drives anti-tumour immunity. Nature 575: 217–223

Castel P, Rauen KA, McCormick F (2020) The duality of human oncoproteins: drivers of cancer and congenital disorders. Nat Rev Cancer 379: 1–15

Chaffer CL, Weinberg RA (2015) How does multistep tumorigenesis really proceed? Cancer Discov 5: 22–24

Chen TH, Wang YH, Wu YH (2011) Developmental exposures to ethanol or dimethylsulfoxide at low concentrations alter locomotor activity in larval zebrafish: implications for behavioral toxicity bioassays. Aquat Toxicol 102: 162–166

Chia W, Somers WG, Wang H (2008) Drosophila neuroblast asymmetric divisions: cell cycle regulators, asymmetric protein localization, and tumorigenesis. J Cell Biol 180: 267–272

Chippalkatti R, Abankwa D (2021) Promotion of cancer cell stemness by Ras. Biochem Soc Trans 49: 467–476

Cirstea IC, Gremer L, Dvorsky R, Zhang SC, Piekorz RP, Zenker M, Ahmadian MR (2013) Diverging gain-of-function mechanisms of two novel KRAS mutations associated with Noonan and cardio-facio-cutaneous syndromes. Hum Mol Genet 22: 262–270

Crespo P, Leon J (2000) Ras proteins in the control of the cell cycle and cell differentiation. Cell Mol Life Sci 57: 1613–1636

de Alvaro C, Martinez N, Rojas JM, Lorenzo M (2005) Sprouty-2 overexpression in C2C12 cells confers myogenic differentiation properties in the presence of FGF2. Mol Biol Cell 16: 4454–4461

Desbordes SC, Placantonakis DG, Ciro A, Socci ND, Lee G, Djaballah H, Studer L (2008) High-throughput screening assay for the identification of compounds regulating self-renewal and differentiation in human embryonic stem cells. Cell Stem Cell 2: 602–612

Dontu G, Abdallah WM, Foley JM, Jackson KW, Clarke MF, Kawamura MJ, Wicha MS (2003) In vitro propagation and transcriptional profiling of human mammary stem/progenitor cells. Genes Dev 17: 1253–1270

Dorman CM, Johnson SE (1999) Activated Raf inhibits avian myogenesis through a MAPK-dependent mechanism. Oncogene 18: 5167–5176

Duggan MC, Regan-Fendt K, Olaverria Salavaggione GN, Howard JH, Stiff AR, Sabella J, Latchana N, Markowitz J, Gru A, Tridandapani S et al (2019) Neuroblastoma RAS viral oncogene homolog mRNA is differentially spliced to give five distinct isoforms: implications for melanoma therapy. Melanoma Res 29: 491–500

Esposito D, Stephen AG, Turbyville TJ, Holderfield M (2019) New weapons to penetrate the armor: Novel reagents and assays developed at the NCI RAS Initiative to enable discovery of RAS therapeutics. Semin Cancer Biol 54: 174–182

Feige P, Brun CE, Ritso M, Rudnicki MA (2018) Orienting Muscle Stem Cells for Regeneration in Homeostasis, Aging, and Disease. Cell Stem Cell 23: 653–664

Fell JB, Fischer JP, Baer BR, Blake JF, Bouhana K, Briere DM, Brown KD, Burgess LE, Burns AC, Burkard MR et al (2020) Identification of the Clinical Development Candidate MRTX849, a Covalent KRAS(G12C) Inhibitor for the Treatment of Cancer. J Med Chem 63: 6679–6693

Garcia-Espana A, Philips MR (2023) Origin and Evolution of RAS Membrane Targeting. Oncogene 42: 1741–1750

Gebregiworgis T, Kano Y, St-Germain J, Radulovich N, Udaskin ML, Mentes A, Huang R, Poon BPK, He W, Valencia-Sama I et al (2021) The Q61H mutation decouples KRAS from upstream regulation and renders cancer cells resistant to SHP2 inhibitors. Nat Commun 12: 6274

Golebiewska A, Brons NH, Bjerkvig R, Niclou SP (2011) Critical appraisal of the side population assay in stem cell and cancer stem cell research. Cell Stem Cell 8: 136–147

Gomez-Lopez S, Lerner RG, Petritsch C (2014) Asymmetric cell division of stem and progenitor cells during homeostasis and cancer. Cell Mol Life Sci 71: 575–597

Goto H, Tomono Y, Ajiro K, Kosako H, Fujita M, Sakurai M, Okawa K, Iwamatsu A, Okigaki T, Takahashi T, Inagaki M (1999) Identification of a novel phosphorylation site on histone H3 coupled with mitotic chromosome condensation. J Biol Chem 274: 25543–25549

Gremer L, Merbitz-Zahradnik T, Dvorsky R, Cirstea IC, Kratz CP, Zenker M, Wittinghofer A, Ahmadian MR (2011) Germline KRAS mutations cause aberrant biochemical and physical properties leading to developmental disorders. Hum Mutat 32: 33–43

Gross AM, Frone M, Gripp KW, Gelb BD, Schoyer L, Schill L, Stronach B, Biesecker LG, Esposito D, Hernandez ER et al (2020) Advancing RAS/RASopathy therapies: An NCI-sponsored intramural and extramural collaboration for the study of RASopathies. Am J Med Genet A 182: 866–876

Gupta PB, Onder TT, Jiang G, Tao K, Kuperwasser C, Weinberg RA, Lander ES (2009) Identification of selective inhibitors of cancer stem cells by high-throughput screening. 138: 645–659

Haghighi F, Dahlmann J, Nakhaei-Rad S, Lang A, Kutschka I, Zenker M, Kensah G, Piekorz RP, Ahmadian MR (2018) bFGF-mediated pluripotency maintenance in human induced pluripotent stem cells is associated with NRAS-MAPK signaling. Cell Commun Signal 16: 96

Hanahan D (2022) Hallmarks of Cancer: New Dimensions. Cancer Discov 12: 31–46

Hatfield I, Harvey I, Yates ER, Redd JR, Reiter LT, Bridges D (2015) The role of TORC1 in muscle development in Drosophila. Sci Rep 5: 9676

Ho AL, Brana I, Haddad R, Bauman J, Bible K, Oosting S, Wong DJ, Ahn MJ, Boni V, Even C et al (2021) Tipifarnib in Head and Neck Squamous Cell Carcinoma With HRAS Mutations. J Clin Oncol 39: 1856–1864

Hobbs GA, Der CJ, Rossman KL (2016) RAS isoforms and mutations in cancer at a glance. J Cell Sci 129: 1287–1292

Hofmann MH, Gmachl M, Ramharter J, Savarese F, Gerlach D, Marszalek JR, Sanderson MP, Kessler D, Trapani F, Arnhof H et al (2021) BI-3406, a Potent and Selective SOS1-KRAS Interaction Inhibitor, Is Effective in KRAS-Driven Cancers through Combined MEK Inhibition. Cancer Discov 11: 142–157

Hood FE, Sahraoui YM, Jenkins RE, Prior IA (2023) Ras protein abundance correlates with Ras isoform mutation patterns in cancer. Oncogene 42: 1224–1232

Hunter JC, Manandhar A, Carrasco MA, Gurbani D, Gondi S, Westover KD (2015) Biochemical and Structural Analysis of Common Cancer-Associated KRAS Mutations. Mol Cancer Res 13: 1325–1335

Janes MR, Zhang J, Li LS, Hansen R, Peters U, Guo X, Chen Y, Babbar A, Firdaus SJ, Darjania L et al (2018) Targeting KRAS Mutant Cancers with a Covalent G12C-Specific Inhibitor. Cell 172: 578–589 e517

Jee J, Jeon H, Hwang D, Sommer P, Park Z, Cechetto J, Dorval T (2012) High content screening for compounds that induce early stages of human embryonic stem cell differentiation. Comb Chem High Throughput Screen 15: 656–665

Jindal GA, Goyal Y, Burdine RD, Rauen KA, Shvartsman SY (2015) RASopathies: unraveling mechanisms with animal models. Dis Model Mech 8: 769–782

Joseph GA, Wang SX, Jacobs CE, Zhou W, Kimble GC, Tse HW, Eash JK, Shavlakadze T, Glass DJ (2019) Partial Inhibition of mTORC1 in Aged Rats Counteracts the Decline in Muscle Mass and Reverses Molecular Signaling Associated with Sarcopenia. Mol Cell Biol 39

Konieczny SF, Drobes BL, Menke SL, Taparowsky EJ (1989) Inhibition of myogenic differentiation by the H-ras oncogene is associated with the down regulation of the MyoD1 gene. Oncogene 4: 473–481

Kovalski JR, Bhaduri A, Zehnder AM, Neela PH, Che Y, Wozniak GG, Khavari PA (2019) The Functional Proximal Proteome of Oncogenic Ras Includes mTORC2. Mol Cell 73: 830–844 e812

Laplante M, Sabatini DM (2013) Regulation of mTORC1 and its impact on gene expression at a glance. J Cell Sci 126: 1713–1719

Lassar AB, Thayer MJ, Overell RW, Weintraub H (1989) Transformation by activated ras or fos prevents myogenesis by inhibiting expression of MyoD1. Cell 58: 659–667

Lau HY, Ramanujulu PM, Guo D, Yang T, Wirawan M, Casey PJ, Go ML, Wang M (2014) An improved isoprenylcysteine carboxylmethyltransferase inhibitor induces cancer cell death and attenuates tumor growth in vivo. Cancer Biol Ther 15: 1280–1291

Lee J, Choi KJ, Lim MJ, Hong F, Choi TG, Tak E, Lee S, Kim YJ, Chang SG, Cho JM et al (2010) Proto-oncogenic H-Ras, K-Ras, and N-Ras are involved in muscle differentiation via phosphatidylinositol 3-kinase. Cell Res 20: 919–934

Li C, Vides A, Kim D, Xue JY, Zhao Y, Lito P (2021) The G protein signaling regulator RGS3 enhances the GTPase activity of KRAS. Science 374: 197–201

Li W, Ma H, Zhang J, Zhu L, Wang C, Yang Y (2017) Unraveling the roles of CD44/CD24 and ALDH1 as cancer stem cell markers in tumorigenesis and metastasis. Sci Rep 7: 13856

Liu J, Kang R, Tang D (2022) The KRAS-G12C inhibitor: activity and resistance. Cancer Gene Ther 29: 875–878

Livak KJ, Schmittgen TD (2001) Analysis of relative gene expression data using real-time quantitative PCR and the 2(-Delta Delta C(T)) Method. Methods 25: 402–408

Maeda Y, Tidyman WE, Ander BP, Pritchard CA, Rauen KA (2021) Ras/MAPK dysregulation in development causes a skeletal myopathy in an activating Braf(L597V) mouse model for cardio-facio-cutaneous syndrome. Dev Dyn 250: 1074–1095

Mathews LA, Keller JM, Goodwin BL, Guha R, Shinn P, Mull R, Thomas CJ, de Kluyver RL, Sayers TJ, Ferrer M (2012) A 1536-well quantitative high-throughput screen to identify compounds targeting cancer stem cells. J Biomol Screen 17: 1231–1242

McDonald ER, 3rd, de Weck A, Schlabach MR, Billy E, Mavrakis KJ, Hoffman GR, Belur D, Castelletti D, Frias E, Gampa K et al (2017) Project DRIVE: A Compendium of Cancer Dependencies and Synthetic Lethal Relationships Uncovered by Large-Scale, Deep RNAi Screening. Cell 170: 577–592 e510

Mo SP, Coulson JM, Prior IA (2018) RAS variant signalling. Biochem Soc Trans 46: 1325–1332

Moore AR, Rosenberg SC, McCormick F, Malek S (2020) RAS-targeted therapies: is the undruggable drugged? Nat Rev Drug Discov 19: 533–552

Morrison SJ, Kimble J (2006) Asymmetric and symmetric stem-cell divisions in development and cancer. Nature 441: 1068–1074

Mounier R, Lantier L, Leclerc J, Sotiropoulos A, Foretz M, Viollet B (2011) Antagonistic control of muscle cell size by AMPK and mTORC1. Cell Cycle 10: 2640–2646

Najumudeen AK, Jaiswal A, Lectez B, Oetken-Lindholm C, Guzman C, Siljamaki E, Posada IM, Lacey E, Aittokallio T, Abankwa D (2016) Cancer stem cell drugs target K-ras signaling in a stemness context. Oncogene 35: 5248–5262

Nassar D, Blanpain C (2016) Cancer Stem Cells: Basic Concepts and Therapeutic Implications. Annu Rev Pathol 11: 47–76

Norris SR, Nunez MF, Verhey KJ (2015) Influence of fluorescent tag on the motility properties of kinesin-1 in single-molecule assays. Biophys J 108: 1133–1143

Okutachi S, Manoharan GB, Kiriazis A, Laurini C, Catillon M, McCormick F, Yli-Kauhaluoma J, Abankwa D (2021) A Covalent Calmodulin Inhibitor as a Tool to Study Cellular Mechanisms of K-Ras-Driven Stemness. Front Cell Dev Biol 9: 665673

Olguin HC, Olwin BB (2004) Pax-7 up-regulation inhibits myogenesis and cell cycle progression in satellite cells: a potential mechanism for self-renewal. Dev Biol 275: 375–388

Olson EN, Spizz G, Tainsky MA (1987) The oncogenic forms of N-ras or H-ras prevent skeletal myoblast differentiation. Mol Cell Biol 7: 2104–2111

Pal R, Mamidi MK, Das AK, Bhonde R (2012) Diverse effects of dimethyl sulfoxide (DMSO) on the differentiation potential of human embryonic stem cells. Arch Toxicol 86: 651–661

Parisi B, Sunnen M, Chippalkatti R, Abankwa DK (2023) A flow-cytometry-based pipeline for the rapid quantification of C2C12 cell differentiation. STAR Protoc 4: 102637

Pavic K, Chippalkatti R, Abankwa D (2022) Drug targeting opportunities en route to Ras nanoclusters. Adv Cancer Res 153: 63–99

Post Y, Clevers H (2019) Defining Adult Stem Cell Function at Its Simplest: The Ability to Replace Lost Cells through Mitosis. Cell Stem Cell 25: 174–183

Prior IA, Hood FE, Hartley JL (2020) The Frequency of Ras Mutations in Cancer. Cancer Res 80: 2969–2974

Punekar SR, Velcheti V, Neel BG, Wong KK (2022) The current state of the art and future trends in RAS-targeted cancer therapies. Nat Rev Clin Oncol 19: 637–655

Quinlan MP, Quatela SE, Philips MR, Settleman J (2008) Activated Kras, but not Hras or Nras, may initiate tumors of endodermal origin via stem cell expansion. Mol Cell Biol 28: 2659–2674

Rabara D, Tran TH, Dharmaiah S, Stephens RM, McCormick F, Simanshu DK, Holderfield M (2019) KRAS G13D sensitivity to neurofibromin-mediated GTP hydrolysis. Proc Natl Acad Sci U S A 116: 22122–22131

Rauen KA (2013) The RASopathies. Annu Rev Genomics Hum Genet 14: 355–369

Rommel C, Clarke BA, Zimmermann S, Nunez L, Rossman R, Reid K, Moelling K, Yancopoulos GD, Glass DJ (1999) Differentiation stage-specific inhibition of the Raf-MEK-ERK pathway by Akt. Science 286: 1738–1741

Scheffzek K, Ahmadian MR, Kabsch W, Wiesmuller L, Lautwein A, Schmitz F, Wittinghofer A (1997) The Ras-RasGAP complex: structural basis for GTPase activation and its loss in oncogenic Ras mutants. Science 277: 333–338

Schmick M, Kraemer A, Bastiaens PI (2015) Ras moves to stay in place. Trends Cell Biol 25: 190–197

Schubbert S, Zenker M, Rowe SL, Boll S, Klein C, Bollag G, van der Burgt I, Musante L, Kalscheuer V, Wehner LE et al (2006) Germline KRAS mutations cause Noonan syndrome. Nat Genet 38: 331–336

She X, Gao Y, Zhao Y, Yin Y, Dong Z (2021) A high-throughput screen identifies inhibitors of lung cancer stem cells. Biomed Pharmacother 140: 111748

Sherman SP, Pyle AD (2013) Small molecule screening with laser cytometry can be used to identify pro-survival molecules in human embryonic stem cells. PLoS One 8: e54948

Shern JF, Chen L, Chmielecki J, Wei JS, Patidar R, Rosenberg M, Ambrogio L, Auclair D, Wang J, Song YK et al (2014) Comprehensive genomic analysis of rhabdomyosarcoma reveals a landscape of alterations affecting a common genetic axis in fusion-positive and fusion-negative tumors. Cancer Discov 4: 216–231

Shu L, Houghton PJ (2009) The mTORC2 complex regulates terminal differentiation of C2C12 myoblasts. Mol Cell Biol 29: 4691–4700

Siddiqui FA, Vukic V, Salminen TA, Abankwa D (2021) Elaiophylin Is a Potent Hsp90/ Cdc37 Protein Interface Inhibitor with K-Ras Nanocluster Selectivity. Biomolecules 11

Siljamaki E, Abankwa D (2016) SPRED1 Interferes with K-ras but Not H-ras Membrane Anchorage and Signaling. Mol Cell Biol 36: 2612–2625

Simanshu DK, Nissley DV, McCormick F (2017) RAS Proteins and Their Regulators in Human Disease. Cell 170: 17–33

Skapek SX, Ferrari A, Gupta AA, Lupo PJ, Butler E, Shipley J, Barr FG, Hawkins DS (2019) Rhabdomyosarcoma. Nat Rev Dis Primers 5: 1

Skoulidis F, Li BT, Dy GK, Price TJ, Falchook GS, Wolf J, Italiano A, Schuler M, Borghaei H, Barlesi F et al (2021) Sotorasib for Lung Cancers with KRAS p.G12C Mutation. N Engl J Med 384: 2371–2381

Steffen CL, Kaya P, Schaffner-Reckinger E, Abankwa D (2023) Eliminating oncogenic RAS: back to the future at the drawing board. Biochem Soc Trans

Stevenson DA, Allen S, Tidyman WE, Carey JC, Viskochil DH, Stevens A, Hanson H, Sheng X, Thompson BA, Okumura MJ et al (2012) Peripheral muscle weakness in RASopathies. Muscle Nerve 46: 394–399

Stevenson DA, Yang FC (2011) The musculoskeletal phenotype of the RASopathies. Am J Med Genet C Semin Med Genet 157C: 90-103

Stowe IB, Mercado EL, Stowe TR, Bell EL, Oses-Prieto JA, Hernandez H, Burlingame AL, McCormick F (2012) A shared molecular mechanism underlies the human rasopathies Legius syndrome and Neurofibromatosis-1. Genes Dev 26: 1421–1426

Tidyman WE, Goodwin AF, Maeda Y, Klein OD, Rauen KA (2022) MEK-inhibitor-mediated rescue of skeletal myopathy caused by activating Hras mutation in a Costello syndrome mouse model. Dis Model Mech 15

Tsai FD, Lopes MS, Zhou M, Court H, Ponce O, Fiordalisi JJ, Gierut JJ, Cox AD, Haigis KM, Philips MR (2015) K-Ras4A splice variant is widely expressed in cancer and uses a hybrid membrane-targeting motif. Proc Natl Acad Sci U S A 112: 779–784

Tsherniak A, Vazquez F, Montgomery PG, Weir BA, Kryukov G, Cowley GS, Gill S, Harrington WF, Pantel S, Krill-Burger JM et al (2017) Defining a Cancer Dependency Map. Cell 170: 564–576 e516

Vasta JD, Peacock DM, Zheng Q, Walker JA, Zhang Z, Zimprich CA, Thomas MR, Beck MT, Binkowski BF, Corona CR et al (2022) KRAS is vulnerable to reversible switch-II pocket engagement in cells. Nat Chem Biol 18: 596–604

Velica P, Bunce CM (2011) A quick, simple and unbiased method to quantify C2C12 myogenic differentiation. Muscle Nerve 44: 366–370

Wakioka T, Sasaki A, Kato R, Shouda T, Matsumoto A, Miyoshi K, Tsuneoka M, Komiya S, Baron R, Yoshimura A (2001) Spred is a Sprouty-related suppressor of Ras signalling. Nature 412: 647–651

Wall VE, Garvey LA, Mehalko JL, Procter LV, Esposito D (2014) Combinatorial assembly of clone libraries using site-specific recombination. Methods Mol Biol 1116: 193–208

Wang MT, Holderfield M, Galeas J, Delrosario R, To MD, Balmain A, McCormick F (2015) K-Ras Promotes Tumorigenicity through Suppression of Non-canonical Wnt Signaling. Cell 163: 1237–1251

Wang X, Allen S, Blake JF, Bowcut V, Briere DM, Calinisan A, Dahlke JR, Fell JB, Fischer JP, Gunn RJ et al (2022) Identification of MRTX1133, a Noncovalent, Potent, and Selective KRAS(G12D) Inhibitor. J Med Chem 65: 3123-3133

Weiswald LB, Bellet D, Dangles-Marie V (2015) Spherical cancer models in tumor biology. Neoplasia 17: 1–15

Weyman CM, Wolfman A (1998) Mitogen-activated protein kinase kinase (MEK) activity is required for inhibition of skeletal muscle differentiation by insulin-like growth factor 1 or fibroblast growth factor 2. Endocrinology 139: 1794–1800

Whyte DB, Kirschmeier P, Hockenberry TN, Nunez-Oliva I, James L, Catino JJ, Bishop WR, Pai JK (1997) K-and N-Ras are geranylgeranylated in cells treated with farnesyl protein transferase inhibitors. J Biol Chem 272: 14459–14464

Xu Q, Wu Z (2000) The insulin-like growth factor-phosphatidylinositol 3-kinase-Akt signaling pathway regulates myogenin expression in normal myogenic cells but not in rhabdomyosarcoma-derived RD cells. J Biol Chem 275: 36750–36757

Yablonka-Reuveni Z, Rivera AJ (1994) Temporal expression of regulatory and structural muscle proteins during myogenesis of satellite cells on isolated adult rat fibers. Dev Biol 164: 588–603

Yan W, Markegard E, Dharmaiah S, Urisman A, Drew M, Esposito D, Scheffzek K, Nissley DV, McCormick F, Simanshu DK (2020) Structural Insights into the SPRED1-Neurofibromin-KRAS

Complex and Disruption of SPRED1-Neurofibromin Interaction by Oncogenic EGFR. Cell Rep 32: 107909

Yelland T, Garcia E, Parry C, Kowalczyk D, Wojnowska M, Gohlke A, Zalar M, Cameron K, Goodwin G, Yu Q et al (2022) Stabilization of the RAS:PDE6D Complex Is a Novel Strategy to Inhibit RAS Signaling. J Med Chem 65: 1898–1914

Yin H, Price F, Rudnicki MA (2013) Satellite cells and the muscle stem cell niche. Physiol Rev 93: 23–67

Yohe ME, Gryder BE, Shern JF, Song YK, Chou HC, Sindiri S, Mendoza A, Patidar R, Zhang X, Guha R et al (2018) MEK inhibition induces MYOG and remodels super-enhancers in RAS-driven rhabdomyosarcoma. Sci Transl Med 10

Yoon MS (2017) mTOR as a Key Regulator in Maintaining Skeletal Muscle Mass. Front Physiol 8: 788

Yoshida N, Yoshida S, Koishi K, Masuda K, Nabeshima Y (1998) Cell heterogeneity upon myogenic differentiation: down-regulation of MyoD and Myf-5 generates ’reserve cells’. J Cell Sci 111 (Pt 6): 769–779

Zembruski NC, Stache V, Haefeli WE, Weiss J (2012) 7-Aminoactinomycin D for apoptosis staining in flow cytometry. Anal Biochem 429: 79–81

