## Supplementary material for "RAS isoform specific activities are disrupted by disease associated mutations during cell differentiation": SI Figures and Tables

**Supplementary Information – Chippalkatti et al.**

**Figure 1 - supplement 1**

(**A,B**) Brightfield images (20 ×) of C2C12 cells with passage numbers P8 (A) and P10 (B) at indicated times in low serum medium. Scale bars, 200 µm.

(**C,D**) Flow cytometric quantification of MyHC+ C2C12 cells with passage number 8 (C) and 10 (D) at indicated times of differentiation; N ≥ 2. Higher passage numbers increase the MyHC+ fraction on day 0 and show a decreased differentiation potential subsequently.

**Figure 3 - supplement 1**

(**A,B**) Representative immunoblots of lysates from C2C12 cells with *KRAS* and *NRAS* (A) and *HRAS* (B) knockdowns on day 0. Cells were differentiated for indicated times in low serum medium; N = 3.

(**C,D**) Quantification of data as in (A,B) for *KRAS* knockdown (C) and effect of *KRAS* siRNA on protein levels of N-Ras and H-Ras to assess specificity (D); N = 3.

(**E-H**) Analogous analysis as in (C,D) was performed for *NRAS* knockdowns (E,F) and *HRAS* knockdowns (G,H); N = 3.

**Figure 3 - supplement 2**

(**A,B**) Quantitative RT-PCR was employed to validate knockdown of indicated *RAS*-transcripts after si*KRAS* treatment of C2C12 cells on day 0. Cells were differentiated for indicated times in low serum medium (A). Analysis of the specificity by quantitating *NRAS*- and *HRAS*-transcripts after the same treatment (B); N = 3, passage number 6-7. Expression levels relative to GAPDH were normalized to samples treated with CTRL siRNA per time point.

(**C-F**) Analogous analysis as in (A,B) was performed for *NRAS* knockdowns (C,D) and *HRAS* knockdowns (E,F); N = 3.

**Figure 3 - supplement 3**

(**A-D**) Representative immunoblots of C2C12 cell lysates (top) and their quantified normalized pERK signals after day 0 knockdown with CTRL siRNA (A) or siRNAs against *KRAS* (B), *NRAS* (C) and *HRAS* (D); N = 3, passage number 6-7.

(**D-F**) Representative immunoblots of C2C12 cell lysates (top) and their quantified normalized pAkt signals after day 0 knockdown with CTRL siRNA (A) or siRNAs against *KRAS* (B), *NRAS* (C) and *HRAS* (D); N = 3, passage number 6-7.

**Figure 4 - supplement 1**

(**A,B**) Flow cytometric analysis of GFP+ fraction of C2C12 cells in comparison to geometric means of intensities for mEGFP-wt K-Ras (A) and mEGFP-K-RasG12V (B) expressing cells; N = 3, passage number 6. Data suggest that differentiation was mainly perturbed by exogenously expressed Ras-constructs on days 1 and 2.

(**C,D**) Flow cytometric analysis of GFP+ fraction of C2C12 cells in 3 distinct GFP-construct expression windows for mEGFP-wt K-Ras (A) and mEGFP-K-RasG12V (B) expressing cells; N = 3.

Relative expression window range (W) with fold signal over background of autofluorescence of non-labelled cells, W1: $10^1$ to $10^2$ RFU (corresponds to GFP low); W2: $10^2$ to $10^3$ RFU; W3: $10^3$ to $10^4$ RFU; W2 + W3 correspond to GFP high.

**Figure 6 - supplement 1**

(**A**) Flow cytometric analysis of MyHC+ C2C12 cells treated with indicated concentrations of DMSO vehicle during the 3-day differentiation; N = 3, passage number 8.

(**B**) Flow cytometric quantification of intact cells based on forward- and side-scatter in samples from (A) indicates treatment induced toxicity if this fraction decreases.

(**C,D**) Flow cytometric analysis of MyHC+ C2C12 cell fractions expressing EGFP-variant tagged wt K-Ras, K-RasG12V or K-RasG12V-S17N on day 3 of differentiation; N = 3, passage number 8. Cells were analyzed in the GFP low (C) and GFP high (D) windows, after transfection on day 0.

(**E-H**) Flow cytometric quantification of intact cells based on forward- and side-scatter of samples as in Figure 7 indicates compound-induced toxicity if this fraction decreases. after treatments with 3 μM AMG 510 (E), ARS-1620 (F), MRTX1257 (G) and MRTX849 (H) expressed by percentages of intact cells; N = 4, passage number 9.

(**I,J**) Flow cytometric quantification of GFP+ 7AAD+ late apoptotic/necrotic C2C12 cells transfected on day 0 with mEGFP-K-RasG12C and with indicated treatments during 3-days of differentiation; N = 3. Cells adherent (I) or in the supernatant (J) were analyzed separately. Statistical significance was calculated using Kruskal-Wallis test with Dunn's post test.

| **Mutation** | **Disease** | **Frequency** | **Intrinsic GTP hydrolysis** | **NF1 GAP-mediated GTP hydrolysis*** | **SOS1-mediated nucleotide exchange** | **RBD binding affinity** |
|---|---|---|---|---|---|---|
| K-Ras G13D | cancer | 12.8 % | impaired | **normal** | **increased** | decreased |
| K-Ras G12V | cancer | 23.1 % | impaired | **impaired** | normal | decreased |
| K-Ras Q61H | cancer | 0.75 % | impaired | **impaired (p120gap)** | mild increase | mild increase (predicted) |
| N-Ras G12V | cancer | 1.7 % | *impaired (predicted based on K-RasG12V)* | ***impaired (predicted)*** | *normal (predicted)* | *decreased (predicted)* |
| H-Ras G12V | cancer | 10.2 % | *impaired (predicted based on K-RasG12V)* | ***impaired*** *(predicted)* | *normal (predicted)* | *decreased (predicted)* |
| K-Ras G12C | cancer | 11.5 % | reduced | **impaired (p120gap)** | normal | mild decrease |
| K-Ras G12D | cancer | 34.1 % | reduced | **impaired** | normal | decreased |

**Table S1. Qualitative changes of biochemical properties of oncogenic Ras mutants.** Information on the biochemical properties was compiled from references given in the main text. *Unless otherwise annotated in the table.

| **Mutation** | **Disease** | **Frequency** | **Intrinsic GTP hydrolysis** | **NF1 GAP-mediated GTP hydrolysis*** | **SOS1-mediated nucleotide exchange** | **RBD binding affinity** |
|---|---|---|---|---|---|---|
| K-Ras T58I | NS | / | reduced | **normal** | normal | decreased |
| K-Ras P34L | NS | / | normal | **impaired** | normal | strongly decreased |
| K-Ras Y71H | NS | / | impaired | **higher** | normal | increased |
| K-Ras D153V | NS | / | reduced | **normal** | normal | normal |
| K-Ras Q22E | CFCS | / | normal | **reduced** | increased | decreased |
| K-Ras V14I | NS | / | normal | **normal** | increased | decreased |
| K-Ras K5E | CS | / | *normal (predicted)* | ***normal*** *(predicted)* | *normal (predicted)* | *normal (predicted)* |
| K-Ras P34R | NS & CFCS | / | normal | **impaired** | normal | strongly decreased |
| K-Ras K5N | CS | / | normal | **normal** | normal | normal |
| K-Ras G60R | CFCS | / | impaired | **impaired** | normal | strongly decreased |

**Table S2. Qualitative changes of biochemical properties of RASopathy K-Ras mutants.** Information on the biochemical properties was compiled from references given in the main text.

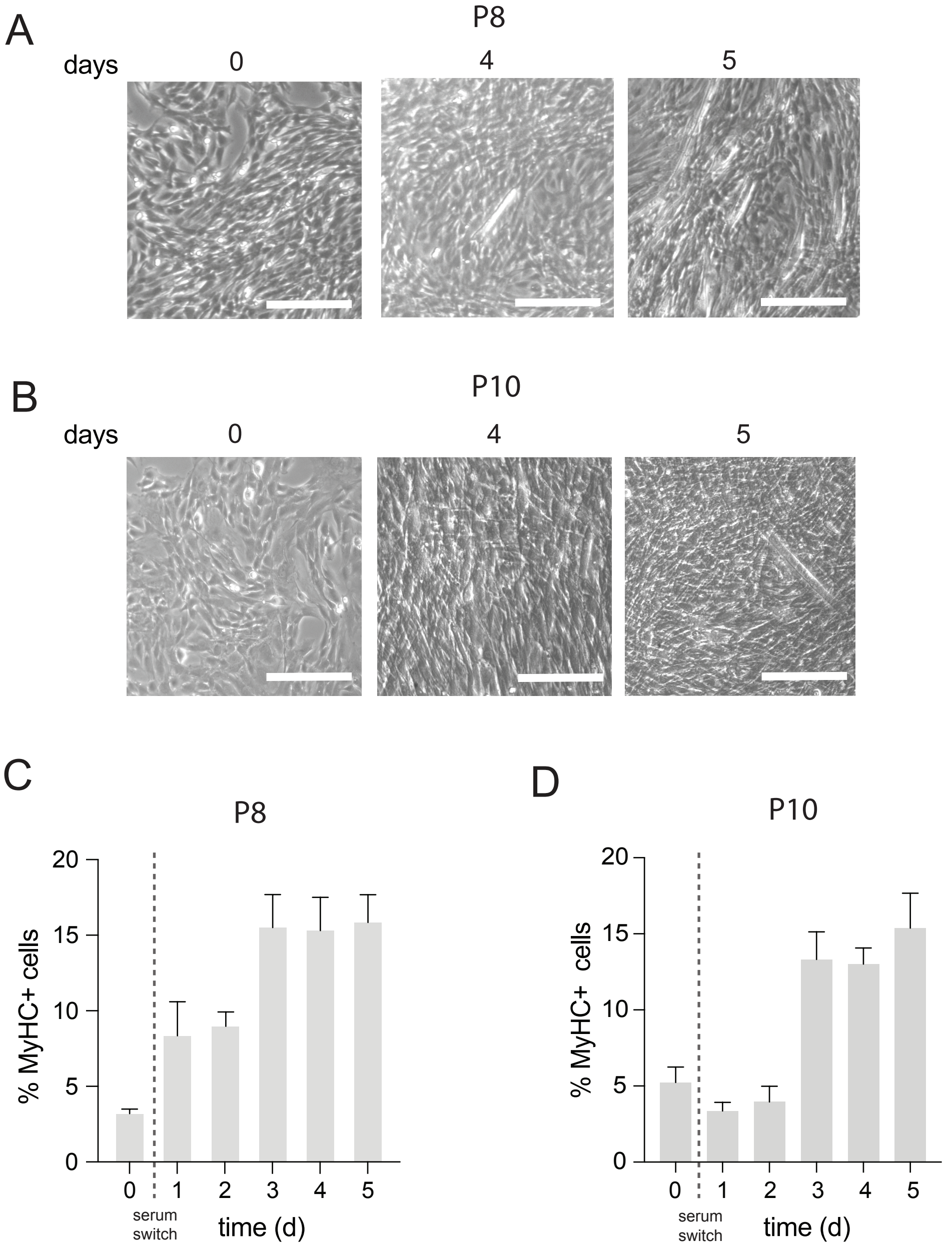


Figure 1
supplement 1

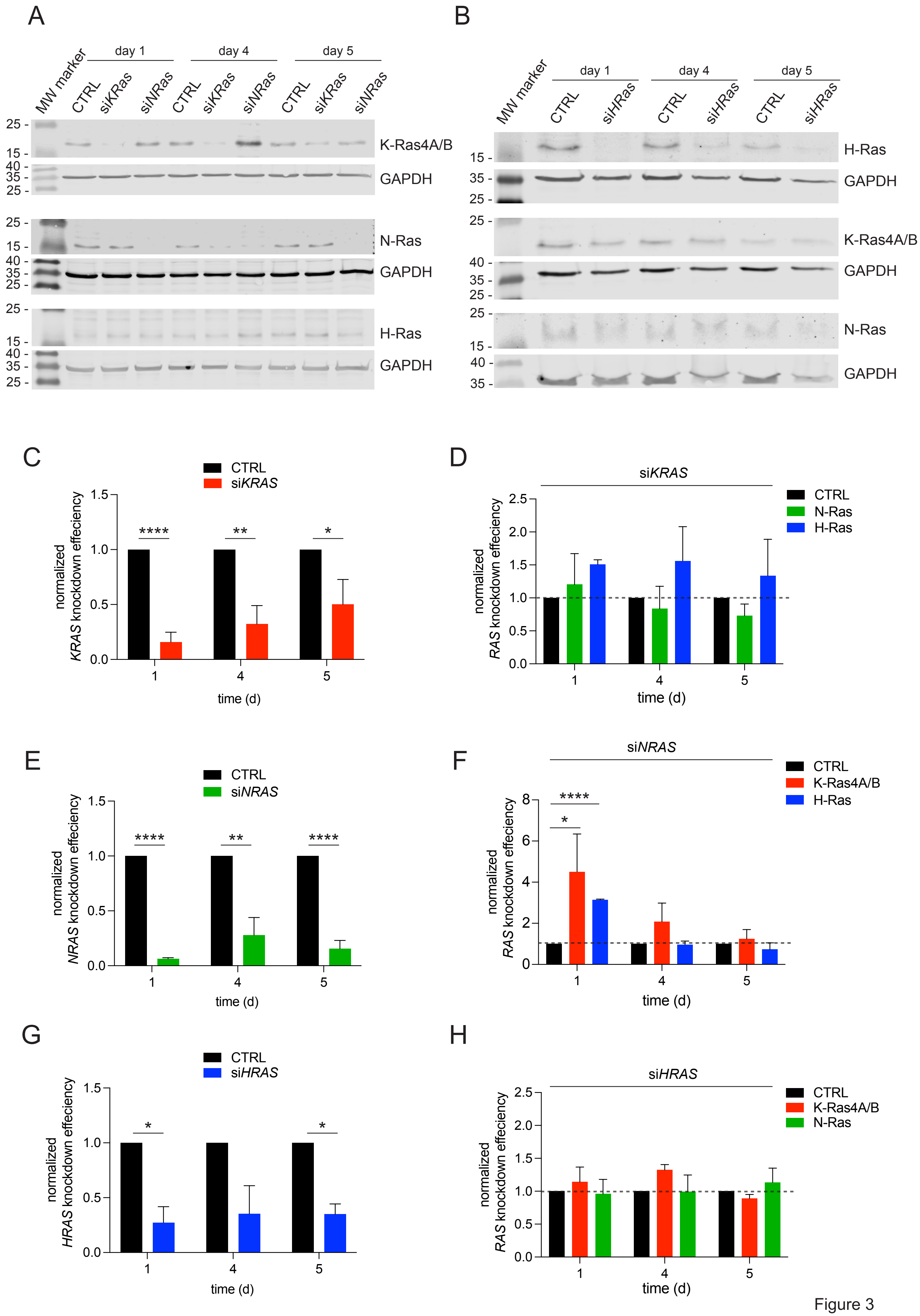


Figure 3
supplement 1

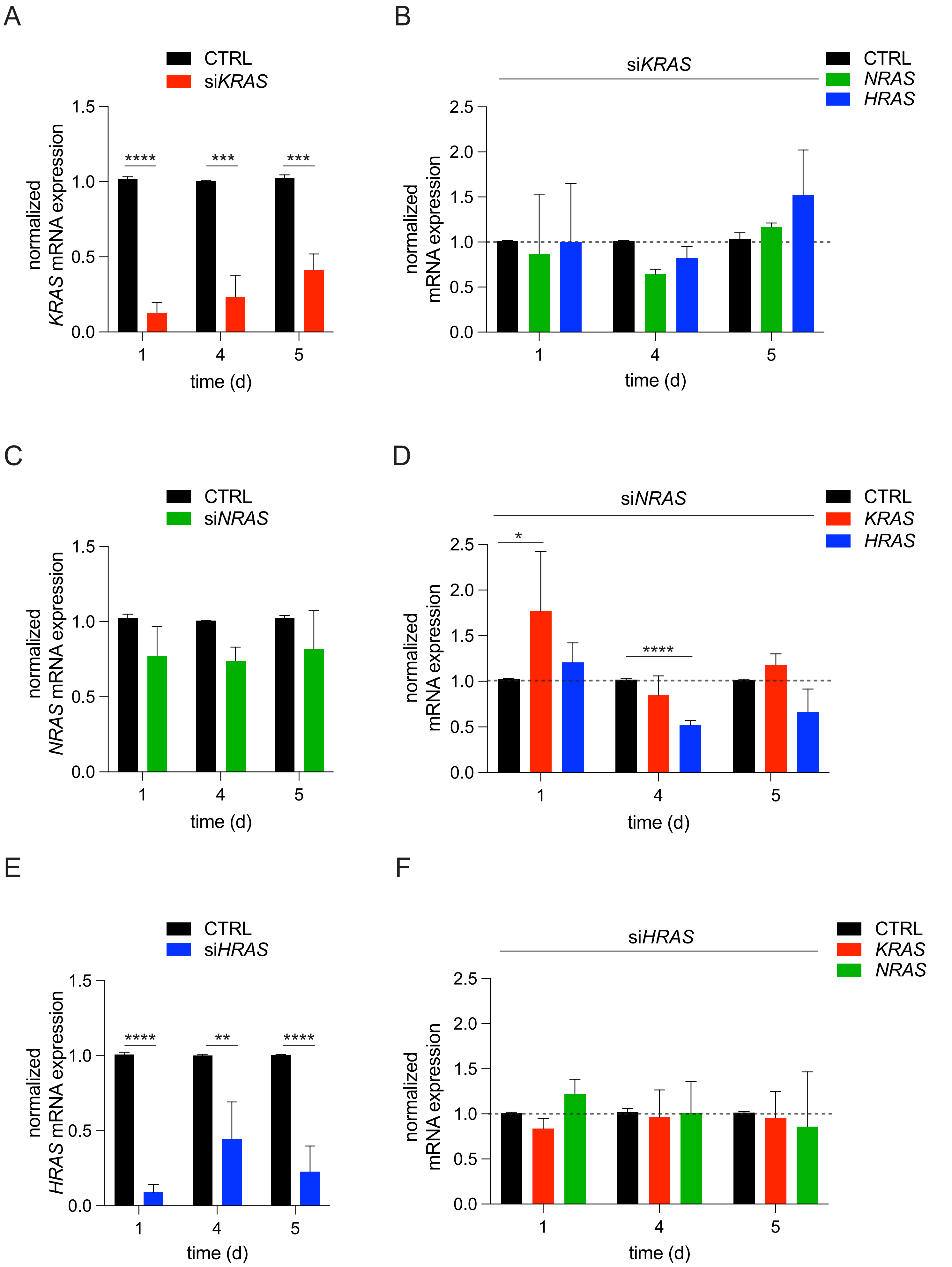


Figure 3
supplement 2

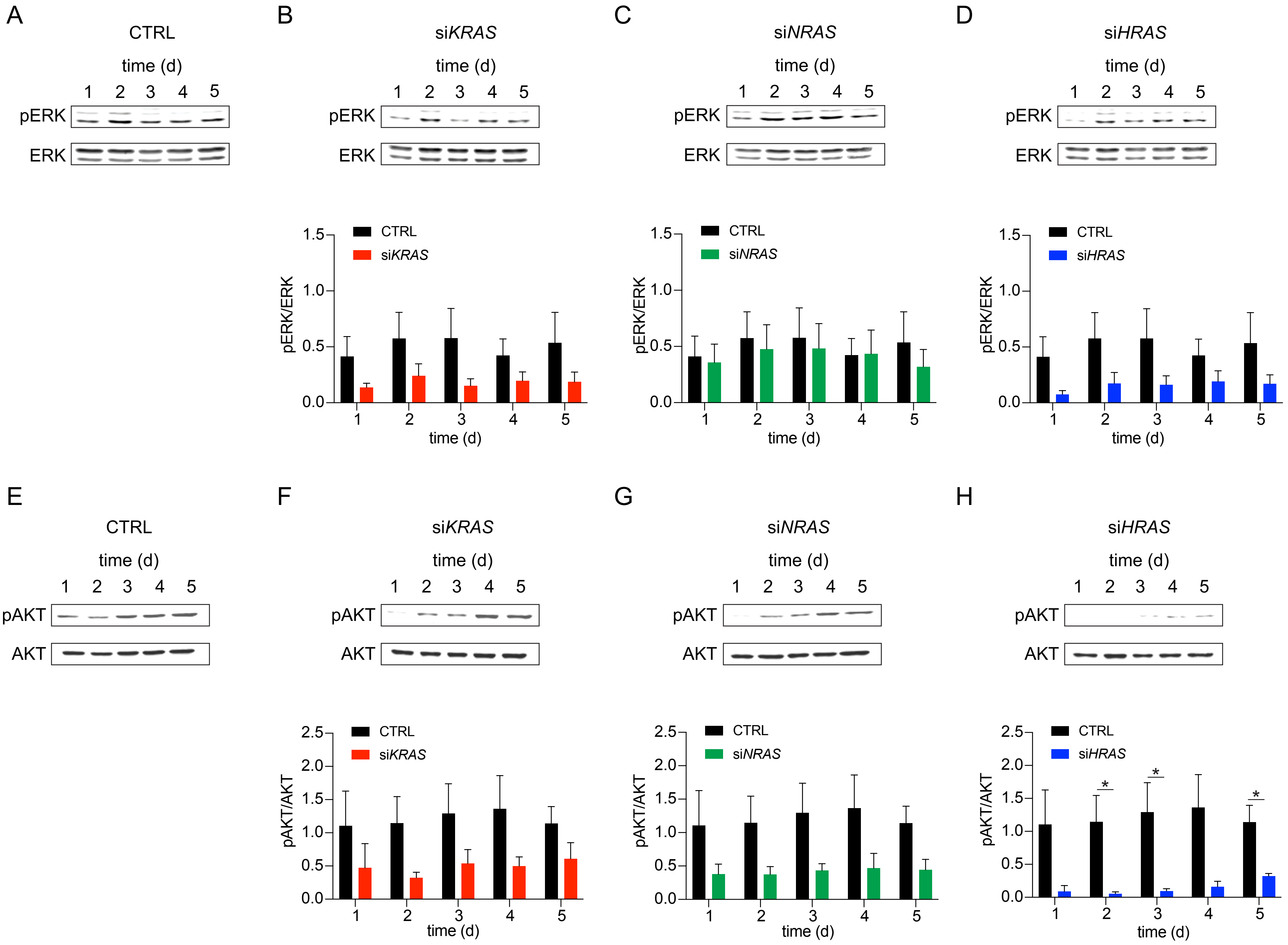


Figure 3
Supplement 3

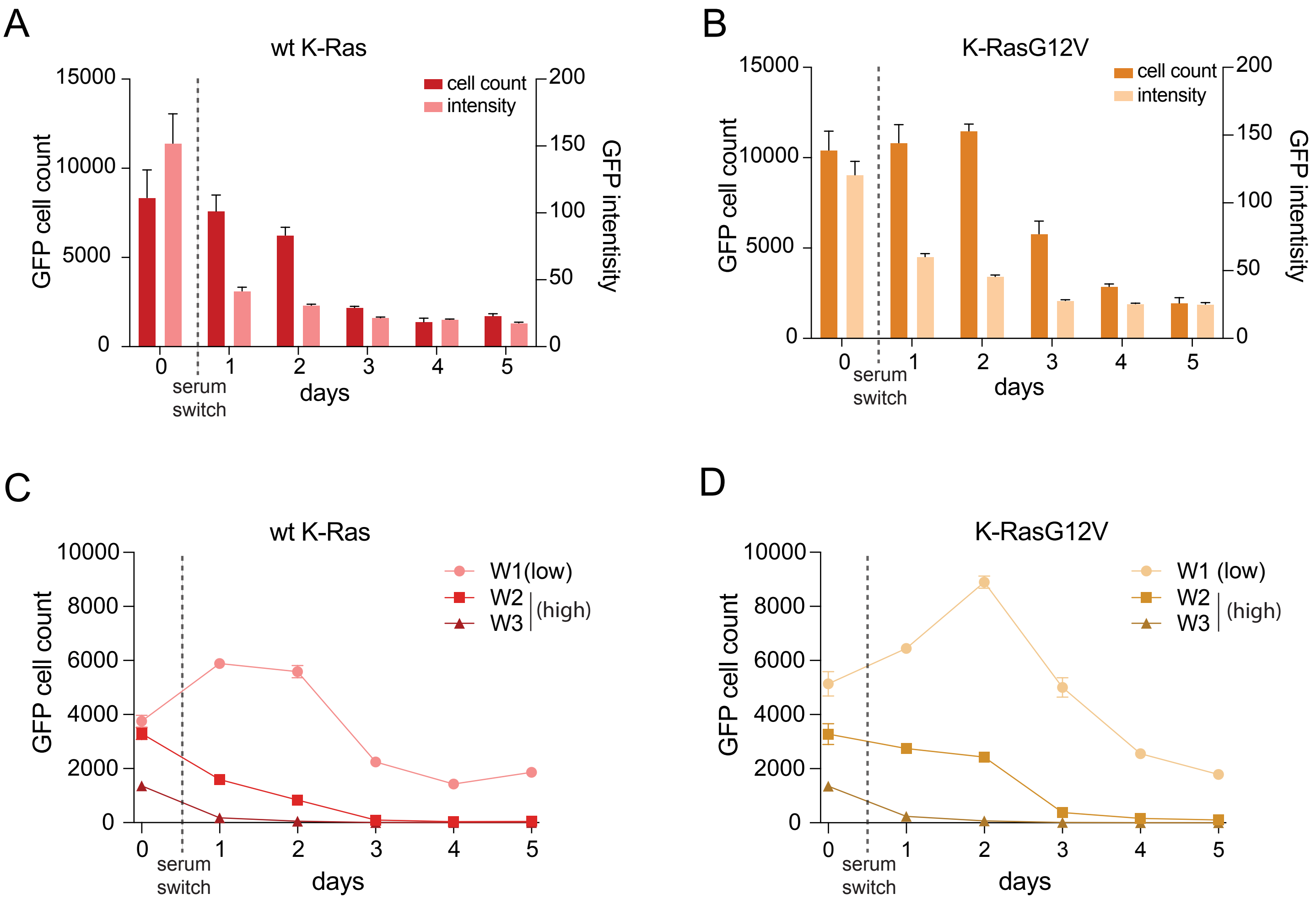


Figure 4
supplement 1

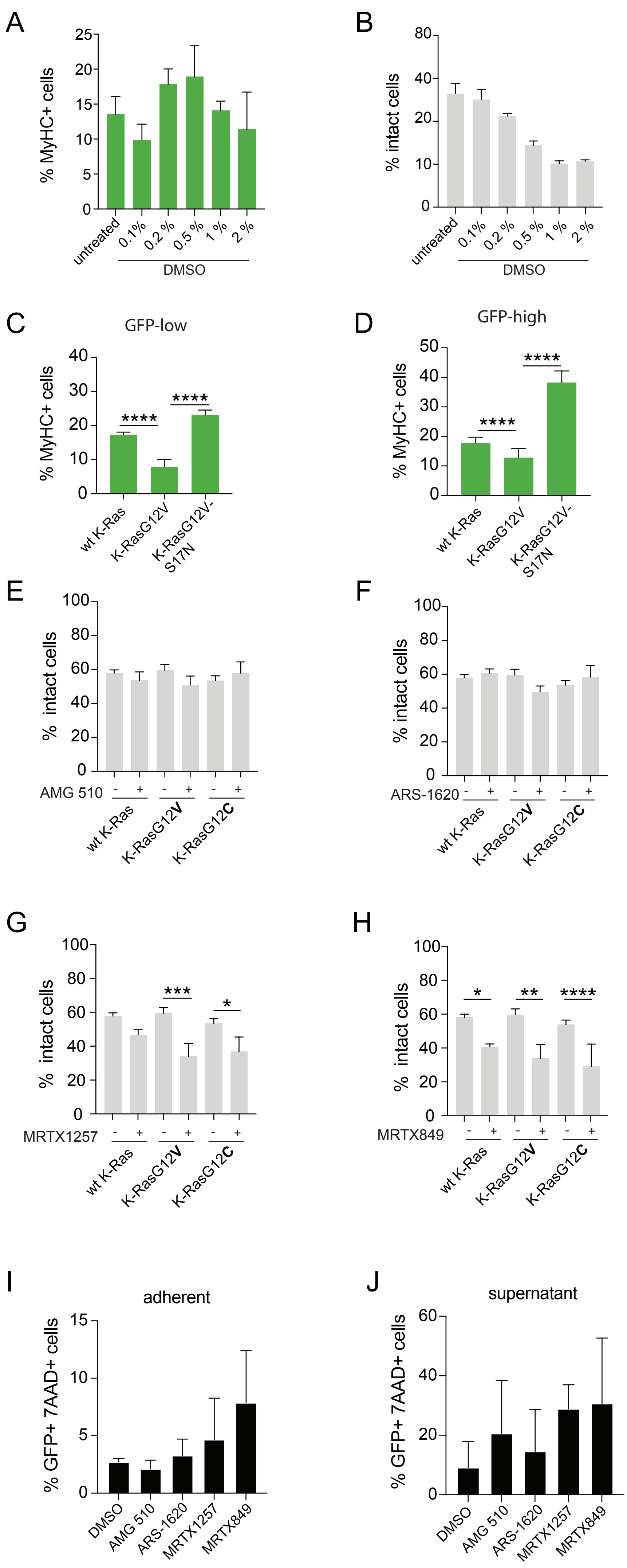


Figure 6 supplement 1
